# Defining the Transcriptional and Epigenetic Basis of Organotypic Endothelial Diversity in the Developing and Adult Mouse

**DOI:** 10.1101/2021.11.15.468651

**Authors:** Manuel E. Cantu Gutierrez, Matthew C. Hill, Gabrielle Largoza, James F. Martin, Joshua D. Wythe

## Abstract

Significant phenotypic differences exist between the vascular endothelium of different organs, including cell-cell junctions, paracellular fluid transport, shape, and mural cell coverage. These organ-specific morphological features ultimately manifest as different functional capacities, as demonstrated by the dramatic differences in capillary permeability between the leaky vessels of the liver compared to the almost impermeable vasculature found in the brain. While these morphological and functional differences have been long appreciated, the molecular basis of endothelial organ specialization remains unclear. To determine the epigenetic and transcriptional mechanisms driving this functional heterogeneity, we profiled accessible chromatin, as well as gene expression, in six different organs, across three distinct time points, during murine development and in adulthood. After identifying both common, and organ-specific DNA motif usage and transcriptional signatures, we then focused our studies on the endothelium of the central nervous system. Using single cell RNA-seq, we identified key gene regulatory networks governing brain blood vessel maturation, including TCF/LEF and FOX transcription factors. Critically, these unique regulatory regions and gene expression signatures are evolutionarily conserved in humans. Collectively, this work provides a valuable resource for identifying the transcriptional regulators controlling organ-specific endothelial specialization and provides novel insight into the gene regulatory networks governing the maturation and maintenance of the cerebrovasculature.

## INTRODUCTION

The endothelium, which lines all blood vessels and is the main component involved in the exchange of nutrients and waste throughout the body, is presumed to have evolved in a common vertebrate ancestor some 500 million years ago, following the divergence of urochordates and cephalochordates (Aird, 2012). Studies in hagfish, the oldest living vertebrate with a closed circulatory system, revealed that the endothelium is molecularly, anatomically, and functionally heterogeneous (Feng et al., 2007; Yano et al., 2007). This suggests that phenotypic heterogeneity is an evolutionarily conserved, core feature of the vascular endothelium. Yet, the molecular basis of this heterogeneity remains poorly understood.

The tubular networks formed by endothelial cells extend throughout the mammalian body, and no cell is more than 100-150 μm away from the capillary vessels, which supply tissues with oxygen and nutrients and also remove cellular waste products (Carmeliet and Jain, 2000). Despite a shared mesodermal origin and a host of common functions, endothelial cells are not a homogenous population (Aird, 2007, 2012; Jambusaria et al., 2020). Indeed, the endothelium varies not only across organs, with diverse physiological functions and anatomical compositions, but also across embryogenesis, allowing vessels to adapt to meet the diverse energetic demands of their surrounding tissues (Kalucka et al., 2020; Marcu et al., 2018; Nolan et al., 2013; Paik et al., 2020). For example, the hepatic sinusoidal capillaries of the liver feature large intercellular gaps (or fenestrae) between endothelial cells and lack an organized basement membrane, which allows for maximal contact and exchange between blood and hepatocytes in the space of Disse (Hwa and Aird, 2007). These fenestrae are essential for receptor-mediated endocytosis of lipoproteins, and allow sinusoidal ECs to function as scavengers, eliminating soluble macromolecular waste. In contrast, the primary function of ECs within the kidney glomeruli is to filter fluids and solutes (Mohamed and Sequeira-Lopez, 2019). While glomerular capillary ECs also possess intercellular fenestrae, these gaps are smaller in glomerular ECs than in their liver sinusoidal counterparts (60-80 nm in diameter vs 100-200 nm). However, glomerular holes in the basement membrane cover more cell surface area (∼20% vs 6-8%, respectively) (Churg and Grishman, 1975). Unlike sinusoidal ECs, glomerular ECs secrete and deposit a glycocalyx, a formidable (60-300 nm thick) cell surface layer of membrane-associated proteoglycans, glycolipids, glycosamines, and associated plasma proteins that forms another filtration barrier (based on charge) (Menzel and Moeller, 2011).

In addition to heterogeneity between organs, ECs *within* organs also display substantial differences. While well-established molecular and functional differences distinguish the endothelium of arterial, arteriole, venous, venule, and capillary vessels (Fish and Wythe, 2015), multiple recent reports have identified additional distinct EC subpopulations within adult mouse organs, such as the lung (Vila Ellis et al., 2020). When one considers the diverse microenvironments within an organ, such as the kidney, where ECs in the vasa recta of the inner medulla exist in a low oxygen, hyperosmolar, hyperkalemic environment, it is perhaps not surprising that a recent study identified up to 24 distinct renal endothelial phenotypes (Dumas et al., 2020). Clearly the adaptations required to thrive in this harsh environment are different than those of capillaries located proximal to alveoli within the oxygen-rich environment of the lung. These diverse functions and phenotypes of ECs demonstrate their inherit phenotypic plasticity, and suggest that cellular heterogeneity is a core property that allows ECs to fulfill their multiple tasks. Conceptually, this makes sense, as the endothelial network that traverses the body must adapt to fulfill the diverse physiological demands of the underlying tissues. In support of this concept, uncoupling endothelial cells from their native microenvironment and local extracellular cues (i.e. cytokines, metabolites, cell-cell contacts with underlying parenchymal cells, etc.) by growing them in culture leads to phenotypic drift, as unique markers and molecular signatures are lost (Aranguren et al., 2013; Burridge and Friedman, 2010; Goldeman et al., 2020; Lacorre et al., 2004). Conversely, *in vivo* transplantation studies showed that the local tissue microenvironment can alter endothelial cell gene expression (Aird et al., 1997).

Despite their residing in distinct locations, endothelium within these various organs all possess the same genome. Thus, their functional diversification likely derives from how the genome is activated via chromatin accessibility and/or epigenetic regulation (Augustin and Koh, 2017; Cleuren et al., 2019). Enhancers, non-coding regions of the genome that modify transcriptional output, are central nodes in transcriptional networks, integrating multiple upstream signals into unified outputs that act to regulate promoter activity and ultimately induce changes in gene expression (Visel et al., 2009b). Several techniques have emerged to map enhancers, which are difficult to predict *a priori* due to their undefined sequence or location (with respect to their target genes). Methods such as immunoprecipitation for unique covalent histone modifications associated with transcriptionally active chromatin (e.g., acetylation of histone H3 lysine 27, H3K27ac) followed by next-generation sequencing (ChIP-seq), or DNase hypersensitivity mapping, have identified potential regulatory elements. However, while most enhancers are DNase hypersensitive, most DNase hypersensitive regions are not active enhancers (Crawford et al., 2006; Thurman et al., 2012). Similarly, while H3K27ac is enriched in cell-type specific enhancers (Crawford et al., 2006; Thurman et al., 2012), this mark alone may not accurately predict enhancers (Dogan et al., 2015). Ep300, a transcriptional co-activator and histone acetyltransferase that catalyzes H3K27 acetylation, is perhaps a stronger indicator of active enhancers (Visel et al., 2009a), yet reproducibility of P300-binding sites has been an issue due to antibody variability (Gasper et al., 2014; Zhou et al., 2017). Additionally, purifying endothelium from different organs for expression profiling or epigenetic studies is not trivial, and complicated FACS procedures represent a serious bottleneck and may introduce artifacts from the time of tissue collection to the time of analysis. Furthermore, the amount of input material required can be daunting if the lineage of interest comprises a small fraction of the cells in a tissue of interest (e.g. the approximately 5,000 endothelial cells of the adult retina, for example). ATAC-seq (Assay for Transposase-Accessible Chromatin using sequencing) overcomes these hurdles, as it uses a robust, transposase enzyme-based method to profile open, accessible chromatin, rather than histone modifications, and requires substantially less input (50,000 nuclei, or less)(Buenrostro et al., 2013).

By combining Cre-dependent expression of a genetically encoded, fluorescently tagged nuclear membrane protein (Sun1-2xsfGFP) (Mo et al., 2015) with an endothelial-specific CreER driver line (Sorensen et al., 2009), we selectively isolated endothelial nuclei from six different organs, across three developmental timepoints, via INTACT (isolation of <u>n</u>uclei <u>ta</u>gged in specific <u>c</u>ell types) (Deal and Henikoff, 2010). As ATAC-Seq requires little biological material (50,000 nuclei), we were able to process the remaining nuclei for transcriptional analysis by RNA-sequencing to define both the shared, and unique, transcriptional and epigenomic features of the vascular endothelium of six different organs during three stages of murine development. Using this strategy, we identified common accessible chromatin regions present in all organs, as well as the DNA-binding motifs within these regions, to define a “core” endothelial transcriptional code involving ETS and SOX family transcription factors. We then mined this data to identity organ-specific, accessible endothelial enhancers in embryonic and postnatal development, as well as in the adult mouse. Analysis of these putative organ-specific, accessible enhancers and promoters revealed transcription factor DNA-binding motifs – which likely govern EC gene expression – within these distinct organs, while gene expression analysis identified the specific transcription factor family member(s) likely driving gene expression through these unique DNA regulatory elements. We extended these observations to examine the transcriptional and epigenetic changes in the vasculature of the central nervous system across developmental time, and through extensive single cell RNA-seq and bioinformatic analysis we identified gene regulatory networks that govern angiogenesis and blood brain barrier maturation in the mouse. Critically, profiling accessible chromatin in human brain endothelial cells determined that the transcriptional networks identified in the mature mouse brain were evolutionarily conserved in humans. Thus, we present a compendium of shared, and unique, transcriptome and epigenetic data across multiple organs, throughout development and adulthood, for identification of the key transcriptional regulators and DNA-binding motifs that govern organ-specific endothelial gene expression of the vascular endothelium.

## RESULTS

### Endothelial Cell Chromatin Accessibility Profiling Using INTACT and ATAC-Seq Across Multiple Organs Over Time

To analyze organ-specific differences in endothelial chromatin accessibility and gene expression, we used a previously validated, endothelial-specific CreERT2 driver line (Cdh5-PAC-CreER) (Sorensen et al., 2009), combined with a Cre-dependent reporter mouse (Rosa26^CAG-lox-stop-lox-Sun1-sfGFP^, denoted as R26^Sun1-sfGFP^) (Mo et al., 2015). Combining these two alleles allows for tissue-specific expression of super folder GFP (sfGFP) in the nuclear envelope of endothelial cells following administration of tamoxifen. This Cre-dependent labeling enabled isolation of <u>n</u>uclei <u>ta</u>gged in specific <u>c</u>ell types (INTACT) via affinity pulldown for sfGFP tagged nuclei (Mo et al., 2015). A mixture of total nuclei was used as a control (i.e. “input”), while Cdh5-CreER-recombined sfGFP-immunoprecipitated nuclei were considered endothelial. Both input and endothelial samples were processed for ATAC-Seq (Buenrostro et al., 2013) and nuclear RNA-seq to identity differentially accessible chromatin and unique transcriptional signatures specific to the endothelium of each different organ (the processing pipeline is shown in Figure 1A). Endothelial cells from the embryonic day 12.5 (E12.5) trunk, brain, and heart, as well as the postnatal day 6 (P6) and adult mouse brain, retina, heart, lung, liver, and kidneys were analyzed (a full list of samples can be found in Supplemental Table 1).

**Figure 1:**
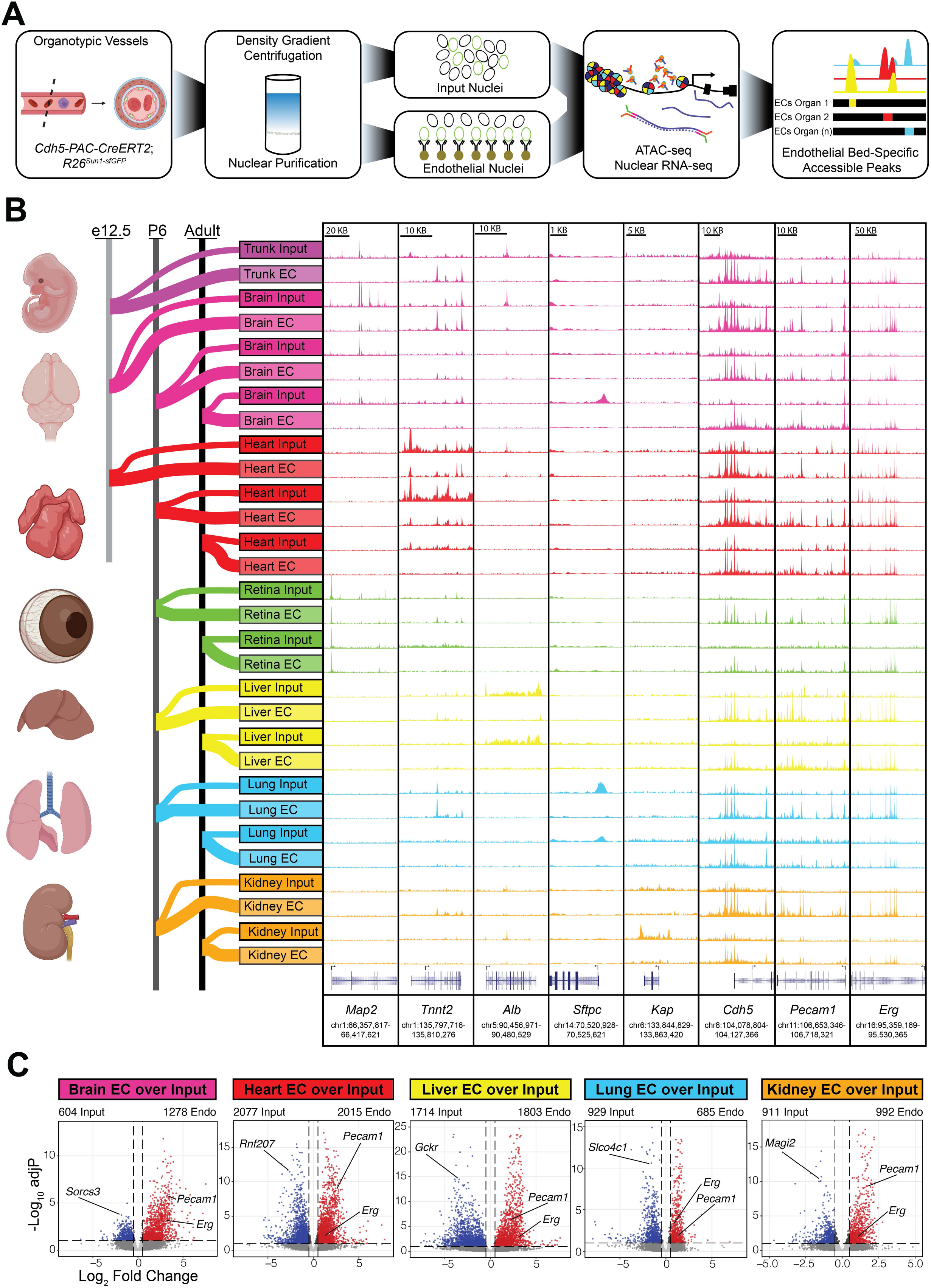
Isolation and Characterization of Tissue-Specific Endothelial Signatures Throughout Development. A) Workflow for genetic affinity tag labelling of ECs using *Cdh5(PAC)-CreERT2* and *R26^Sun1-sfGFP^* mice for isolation of nuclei tagged in specific cell types (INTACT). (Far left) Representative schematic of a blood vessel with GFP-tagged nuclei. Nuclear isolation was followed by RNA-seq profiling of nuclear transcripts and ATAC-seq mapping of accessible chromatin and aligning reads to the mouse genome (far right). B) (Far left) Various tissues and time points used to map endothelial cell diversity in the developing (E12.5), postnatal (P6) and adult (2 months of age) mouse. (Far right) Representative genome browser tracks from ATAC-seq highlight accessible chromatin regions unique to organ-specific genes like *Map2* in neurons, *Tnnt2* in cardiomyocytes, *Alb* in hepatocytes, *Sftpc* in alveolar cells of the lung, and *Kap* in proximal tubule cells of the kidney, and endothelial cells including *Cdh5*, *Pecam1* and *Erg*. C) Volcano plots show differentially expressed genes between the endothelium (red) and input nuclei (blue). All developmental timepoints (E12.5-Adult) are combined and treated as a single timepoint for these analyses.

To confirm the integrity of our organ collection and tissue processing pipeline, we analyzed the chromatin accessibility for genomic loci whose transcripts are enriched in the non-EC major cellular constituents of each organ sampled (i.e. neurons in the brain, cardiomyocytes in the heart, etc.). Accordingly, Map2 (Microtubule Associated Protein 2) (Kanai and Hirokawa, 1995; Matus et al., 1981) accessibility was enriched in brain input comparted to EC nuclei, while Tnnt2 (Troponin T2, Cardiac) (Wang et al., 2001; Yan et al., 2016) was elevated in the heart input, Alb (Albumin) (Kimball et al., 1995; Redman, 1969) in the liver input, Sftpc (Surfactant pulmonary associated protein C) (Nureki et al., 2018) was elevated in in the lung input, and open chromatin surrounding the Kap (Kidney androgen-regulated protein) locus was enriched in the kidney input (Toole et al., 1979). Next, we verified that pan-vascular markers, such as Cdh5 (encoding VE-Cadherin) (Harris and Nelson, 2010), Pecam1 (CD31) (Newman, 1994) and Erg (ERG) (Birdsey et al., 2008) featured increased chromatin accessibility in isolated EC nuclei compared to total input across all tissue types and timepoints (Figures 1B). Examination of our nuclear RNA-seq results confirmed the purity of each organ isolation, as well as the selective enrichment of endothelial nuclei over total input. For example, the neuronal synaptic receptor Sorcs3 (sortilin-related receptor CNS expressed 3) was enriched in the brain (Christiansen et al., 2017), while ubiquitin ligase Rnf207 (RING finger protein 207) was differentially expressed in the heart (Roder et al., 2014), Gckr (Glucokinase regulatory protein) in the liver (Wang et al., 2013), Slco4c1 (Solute carrier organic anion transporter family, member 4C1) in the lung (Leikauf et al., 2012), and Magi-2 (MAGUK Inverted 2) in the kidney (Balbas et al., 2014), yet these transcripts were depleted in the endothelial nuclei of each organ, respectively. Conversely, the EC-enriched transcripts Pecam1 and Erg (Ets Related Gene) were enriched in all endothelial nuclei samples, confirming the specificity of our experimental approach (Figure 1C).

### Endothelial Cells Feature a Core Epigenetic Landscape Across Time and Space

After confirming the integrity of our processing pipeline, we next investigated whether endothelium from different organs and at unique developmental stages share a common “core” of accessible chromatin regions and a shared transcriptional signature. We identified 2,646 endothelial-enriched accessible regions common to the endothelium of all organs (Figure 2A, Supplemental Table 1). As non-coding regions typically lack annotated biological function, we used the Genomic Regions Enrichment of Annotations Tool (GREAT) (McLean et al., 2010) to computationally identify genes associated with these open chromatin regions, and then queried these genes for shared functions using gene ontology (GO) analysis. Vascular development, blood vessel morphogenesis, and angiogenesis were among the top GO terms common to endothelia across all organs (Figure 2B, 2C). If these accessible regions function as putative enhancers, or represent accessible proximal promoters, we hypothesized that the transcription factor motifs present in these core, common gene regulatory regions might play an important role in endothelial cell biology. To investigate this, Hypergeometric Optimization of Motif EnRichment (HOMER) (Heinz et al., 2010) was used to identify transcription factor motifs enriched in these accessible regions. The ETS family of transcription factors, including ERG and FLI1 (Friend Leukemia Integration 1), are crucial for endothelial development (Abedin et al., 2014; Fish et al., 2017; Vijayaraj et al., 2012; Wythe et al., 2013) and were the most significantly enriched motifs in these commonly accessible regions (or peaks) (Figure 2D). Notably, motifs for the ETS family members ETV2 (ETS Variant Transcription Factor 2, also known as ER71) and ETV1 (ETS Variant Transcription Factor 1) were also significantly enriched, but their transcripts were not detected by RNA-seq (data not shown). Previously, an ETS-dependent enhancer within intron three of *Delta Like 4* (*Dll4*) – regulated by the ETS family member ERG (Wythe et al., 2013) – as well as an upstream enhancer in Endoglin (*Eng*) – regulated by the ETS factors FLI1, ERG and ELF1 (E74-like factor 1) – were validated *in vivo* (Pimanda et al., 2006). These same ETS-dependent enhancers were identified by our analyses (Figure 2E, F). Motifs for the SOX (SRY related-HMG box) family of transcription factors were the second most abundant known DNA binding sites present in regions of open chromatin within the endothelium (Figure 2D). The SOXF subfamily (*Sox7*, *17*, and *18*) shows partial redundancy in controlling angiogenesis and vascular maintenance (Chiang et al., 2017; Lee et al., 2014; Zhou et al., 2015), and *Sox17* was previously shown to regulate arterial differentiation in mice (Corada et al., 2013) and to control endothelial to hematopoietic transition (Lizama et al., 2015). Moreover, the SOXB1 subfamily member *Sox2* has also been implicated in endothelial differentiation *in vitro* (Yao et al., 2019b) and in cerebral arteriovenous malformation *in vivo* (Yao et al., 2019a). Finally, motifs for the Forkhead Box (FOX) family member FOXO1, which regulates angiogenesis and endothelial senescence and metabolism (Paik et al., 2007; Potente et al., 2005; Rudnicki et al., 2018; Wilhelm et al., 2016), were also enriched across all organs.

**Figure 2:**
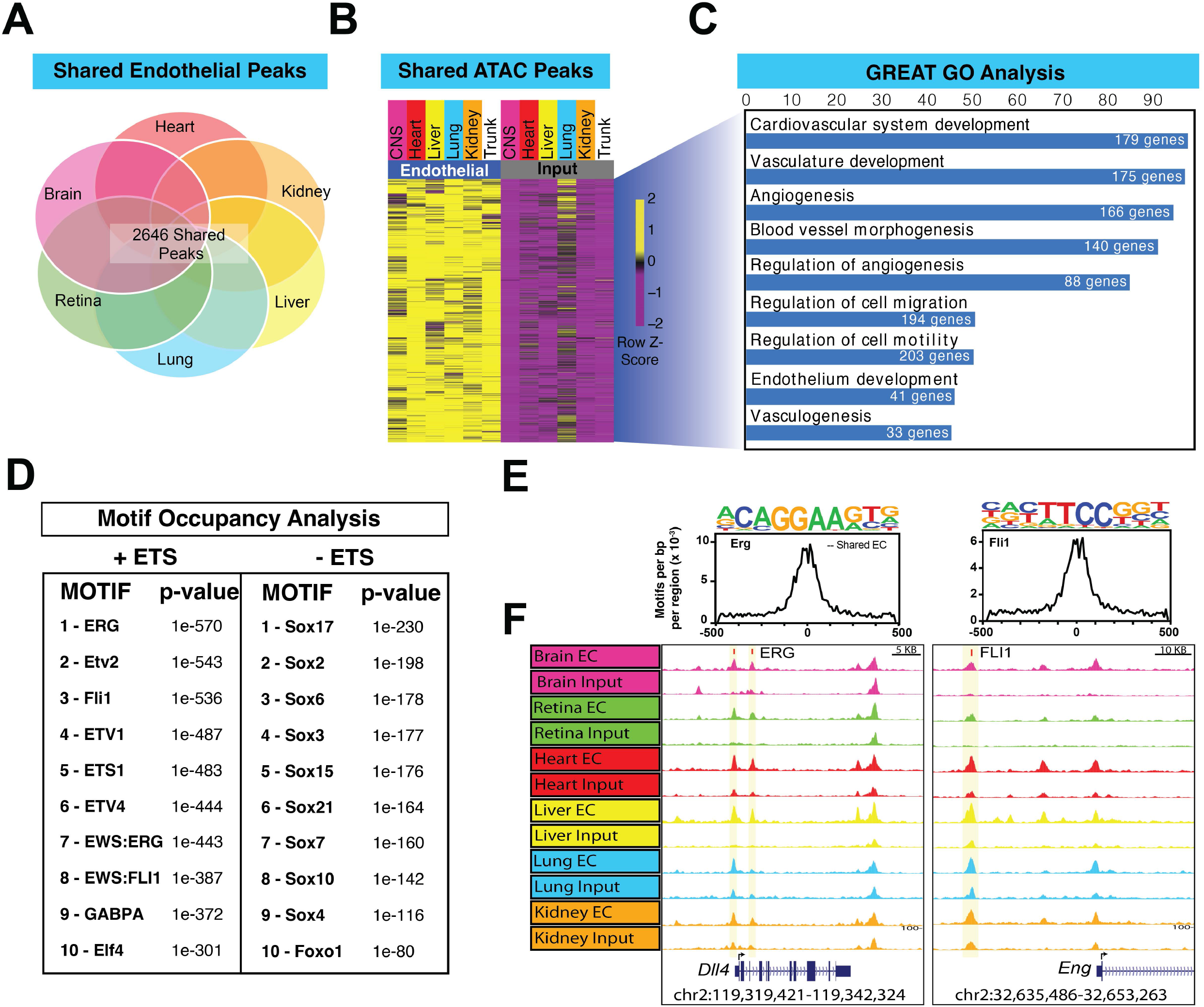
Endothelial Cells from Diverse Organs Share a Core Epigenetic Signature. A) Venn diagram showing the overlap of open chromatin regions (2,646 peaks) between murine heart, kidney, liver, lung, retina, and brain endothelium. B) Heatmap of shared peaks across the endothelial and input datasets. C) GREAT analysis of common peaks showing gene ontology terms related to cardiovascular development and angiogenesis, among others. D) Top 20 transcription factor DNA binding motifs in shared peaks along with their p-value as determined using HOMER. E) Top, position weight matrix (PWM) for transcription factor DNA binding sites, with the inset box showing the frequency of motif occurrence as distance from the center of the peak within accessible DNA regions as determined by ATAC-seq. F) Representative genome browser tracks from ATAC-seq data highlighting accessible chromatin regions in the adult endothelium and representative DNA binding sites (red rectangle) for the transcription factors identified in panel F for the *Delta Like 4* (*Dll4*) and *Endoglin* (*Eng*) loci.

### Organ-Enriched Regions of Accessible Chromatin and Unique Transcription Factor Motifs Across the Endothelium

After characterizing uniformly accessible chromatin regions within the endothelium, and the potential transcription factors that act upon them, we focused our efforts on identifying organ-enriched, endothelial-specific epigenetic signatures from the remaining 90,112peaks. Merging the three timepoints (E12.5, P6.5, and Adult) of each organ to a single dataset, we identified 45,075 EC-enriched peaks that showed differential chromatin accessibility across organs (Figure 3A, Supplemental Table 2). As the brain and retina are both central nervous system (CNS)-derived organs, their data were merged and compared to all other individual organs. We identified 6,550 peaks unique to the CNS vasculature; 11,302 regions specific to the endothelia of the heart; 9,102 to the vessels within the liver; 2,102 open regions in the lung endothelium; and 3,360 peaks in the kidney vasculature (Figure 3A). GREAT (McLean et al., 2010) was used to annotate these regions to nearby genes, and the linked genes were then filtered for enriched gene expression in the endothelium using our nuclear RNA-sequencing data (qvalue < 0.1 and log2Fold change > 0.5). This final list of genes was then used to identify GO terms enriched in each organ (Figure 3B). Brain-enriched regions of open chromatin in the endothelium were associated with genes related to the WNT signaling pathway, as well as cell-cell signaling regulated by WNT. The liver vasculature featured enriched GO terms in the categories of protein phosphorylation and cell adhesion, while the lung endothelium featured enriched terms such as circulatory system processes. The vasculature of the heart and kidney showed enrichment of genes related to semaphorin-plexin signaling, while the heart also showed enrichment for the Notch signaling pathway.

**Figure 3:**
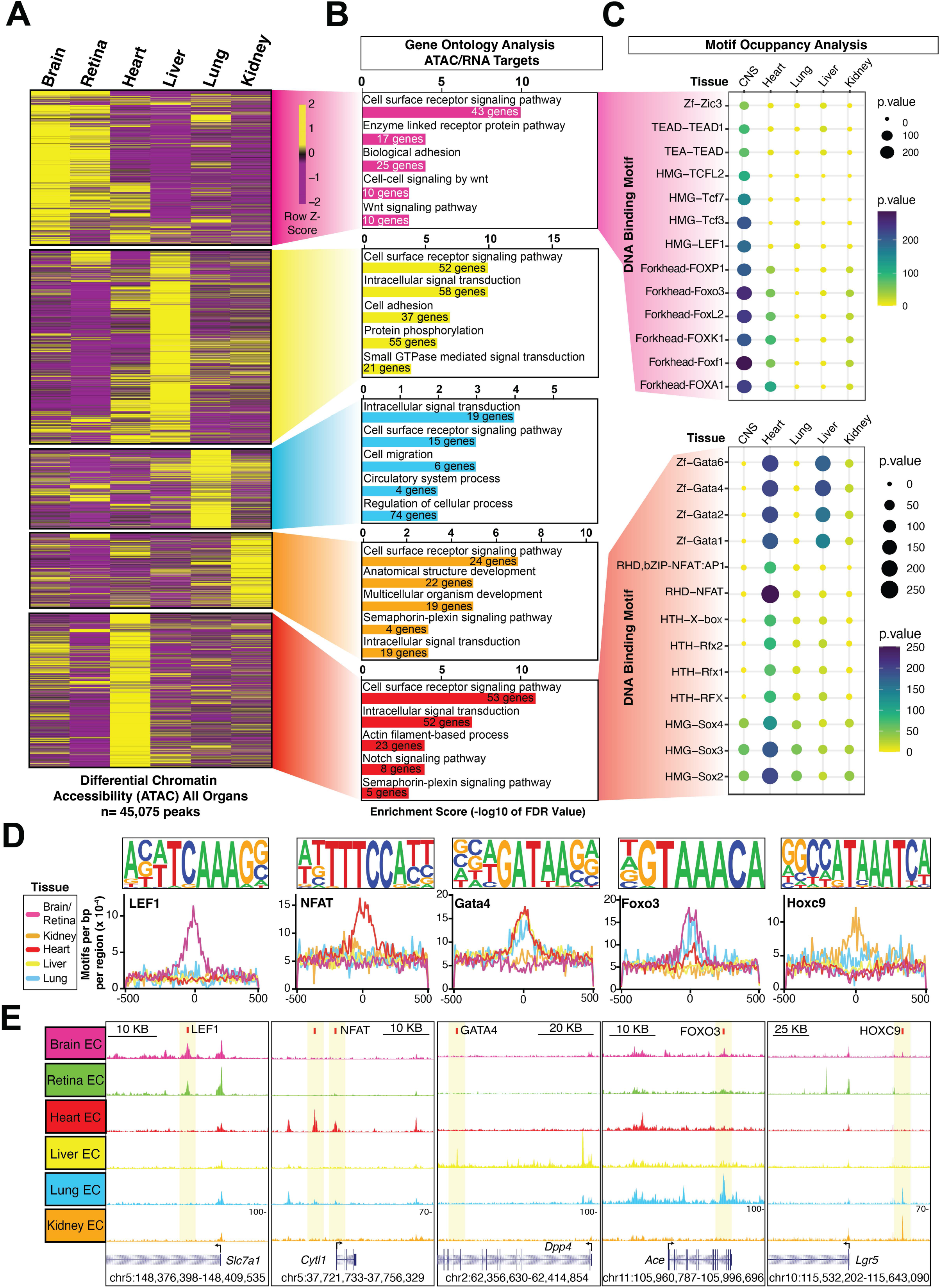
Profiling Accessible Chromatin and Expressed Transcripts Identifies Organ-Specific Endothelial Signatures. A) A heatmap shows differentially accessible regions of open chromatin in the murine brain, retina, heart, liver, lung, and kidney endothelium (45,075 peaks) identified by ATAC-seq. B) Top biological processes from GREAT analysis across differentially accessible peaks in each organ. Only regions annotated to endothelial enriched genes (determined by RNA-sequencing) were used in the analysis. C) Top transcription factor motifs in regions of open chromatin in the brain and heart. D) Enriched motifs found by HOMER in each organ. Position weight matrix (PWM) shown over frequency of motif as distance from peak center. E) Representative genome browser tracks from ATAC-seq highlighting accessible organ-specific chromatin regions in endothelial-enriched transcripts.

Next, to determine which transcription factors recognize (and potentially occupy) these regions of open chromatin in the vessels of each specific tissue, we compared motif occupancy across all organs (Figure 3C, Supplemental Figure 1). In the brain and retina, canonical WNT signaling pathway-related factors play an essential role in the development of the blood brain barrier (Daneman et al., 2009; Hupe et al., 2017; Liebner et al., 2008; Stenman et al., 2008). Among the canonical WNT signaling-related transcription factors found, motifs for ZIC3, TCF3, TCF4, TCF7 and LEF1 were preferentially enriched in the brain endothelium compared to other organs (Figure 3C, Supplemental Figure 1). Additionally, DNA binding motifs for FOX transcription factors were also overrepresented in the brain. To our knowledge, roles for FOXP1, FOXK1, FOXF1 and FOXA1 have not been reported in blood brain barrier development. However, expression of *Foxo3* in the CNS was shown previously, where its downregulation was reported to ameliorate brain damage after cerebral hemorrhage (Xie et al., 2021), and *Foxl2* transcripts are reportedly enriched in the brain endothelium (Hupe et al., 2017).

The heart and liver shared motifs for members of the zinc family of transcription factors GATA1, GATA2, GATA4 and GATA6 (Figure 3C, Supplemental Figure 1). GATA1 has been described as a potential regulator of endothelial cell function in the heart and liver (Fan et al., 2009). GATA2, a master regulator of primitive and definitive hematopoiesis in the liver (de Pater et al., 2013; Lim et al., 2012), is required for endothelial to hematopoietic transition (EHT) and vascular integrity in mice, and promotes the generation of hemogenic endothelial progenitors and represses induction of cardiomyocyte-related genes from human mesoderm (Castano et al., 2019). GATA4 is required for heart valve development (Rivera-Feliciano et al., 2006) and atrial septum formation (Nadeau et al., 2010). In the liver, GATA4 controls the development of liver sinusoidal endothelium (Geraud et al., 2017), while GATA6 is involved in cardiovascular morphogenesis (Lepore et al., 2006) and liver development (Zhao et al., 2005).

Motifs for nuclear factor of activated T cells (NFAT) transcription factors were specifically enriched in the endothelium of the heart (Figure 3C, Supplemental Figure 1). *NFATc* genes (*NFATc1-c4*) play key roles in cardiac morphogenesis. *Nfatc1* is a canonical marker of the endocardium and is required for normal cardiac valve and septal morphogenesis (de la Pompa et al., 1998; Ranger et al., 1998), as well as coronary vessel angiogenesis (Zeini et al., 2009), while *Nfatc3/c4* null embryos, and mutants for their upstream regulator in the heart *Calcineurin* (*Cnb1*), both die at E11.5 with excessive vascular growth (Graef et al., 2001). Motifs for helix-turn-helix (HTH) and winged helix Regulatory Factor binding to the X-box (RFXs) are also enriched in the heart (Figure 3A-C) (Sugiaman-Trapman et al., 2018). Of these enriched motifs, only HTH-X-box is involved in heart (Duan et al., 2016), as a role for DNA-binding *Regulatory Factor 1* and *2* (*Rfx1*, *Rfx2*) in the heart has not been shown.

While SOX2, SOX3 and SOX4 motifs were moderately enriched in endothelium across all organs, they were particularly enriched in the heart (Figure 3C, Supplemental Figure 1). To our knowledge, a role for *Sox2* and *Sox3* in the cardiac vascular endothelium or endocardium has yet to be shown. However, *Sox4* is required for outflow tract morphogenesis (Schilham et al., 1996) and controls *Tbx3* expression in the endocardium (Boogerd et al., 2011). LEF1, NFAT and HOXC9 motifs were enriched in the brain, heart, and kidney, while GATA4 was over-represented in the lung, liver, and heart, and FOXO3 motifs were increased in the brain, and heart (Figure 3A-C).

Notably, motifs for the large MAF (musculoaponeurotic fibrosarcoma) basic leucine zipper (bZip transcription factors) MAFA and MAFB were enriched in the liver endothelium (Figure 3C, Supplemental Figure 1). MAF transcription factors are known to interact with ETS1 or SOX TFs in promoter and enhancer modules (Yang and Cvekl, 2007). MAFb is involved in endothelial sprouting during angiogenesis (Jeong et al., 2017) and lymphangiogenesis (Dieterich et al., 2020). A third member of the large MAF family, c-MAF, was not present in our motif analysis but it has been directly involved in liver sinusoidal endothelial cell marker induction (de Haan et al., 2020).

Importantly, the aforementioned DNA binding motifs were usually enriched in the center of regions of open chromatin for each organ (Figure 3D, Supplemental Table 3), suggesting these factors may be driving chromatin accessibility via acting as pioneer factors or functioning as transcriptional enhancers. Several of these accessible regions and DNA binding motifs occurred within, or nearby, loci of transcripts that are elevated in these individual organs (Figure 3E). For example, *Solute Carrier Family 7 member 1* (*Slc7a1*), which encodes a cationic amino acid transporter that is enriched the endothelium of the mature brain (Nalecz, 2017; Zaragoza, 2020), contains a unique region of open chromatin downstream from the TSS that is unique to the CNS endothelium, and this region contains a LEF1 motif (Figure 3E). *Cytokine-like 1* (*Cytl1*), a novel endocardial gene (Feng et al., 2019), contained four regions of open chromatin unique to the cardiac endothelium, two of which possessed an NFAT motif. *Dipeptidyl-peptidase 4* (*Dpp4*), which encodes a serine protease secreted within the liver endothelium and hepatocytes (Varin et al., 2019), has a liver endothelial specific region of open chromatin that contains a GATA4 motif. *Angiotensin-converting enzyme* (*Ace*), expressed throughout the endothelium, contains a lung specific intronic region of open chromatin with a FOXO3 motif. Finally, the WNT pathway co-receptor, *Leucine-rich repeat-containing G-protein coupled receptor 5* (*Lgr5*) (Wilson et al., 2020), features a kidney-specific region of open chromatin upstream of its promoter with a HOXC9 motif.

### Maturation Specific Regions of Accessible Chromatin and Unique Transcription Factor Motifs in the Developing and Adult CNS Endothelium

After defining the global changes in chromatin accessibility across all organs, we next examined how chromatin organization in the endothelium of each organ varied during development. Focusing on the CNS, we identified 22,182 peaks from E12.5, P6 and Adult (2-month-old) endothelium specific to the brain or overlapping between the brain and retina. After annotating peaks to nearby genes using GREAT (McLean et al., 2010), we then filtered these data for those genes whose transcripts were enriched in the endothelium compared to input (qvalue < 0.1 and log2Fold change > 0.5). These targets were then used for Gene Ontology analysis of biological function (FDR < 0.05) (Figure 4A, B, Supplemental Table 4).

**Figure 4:**
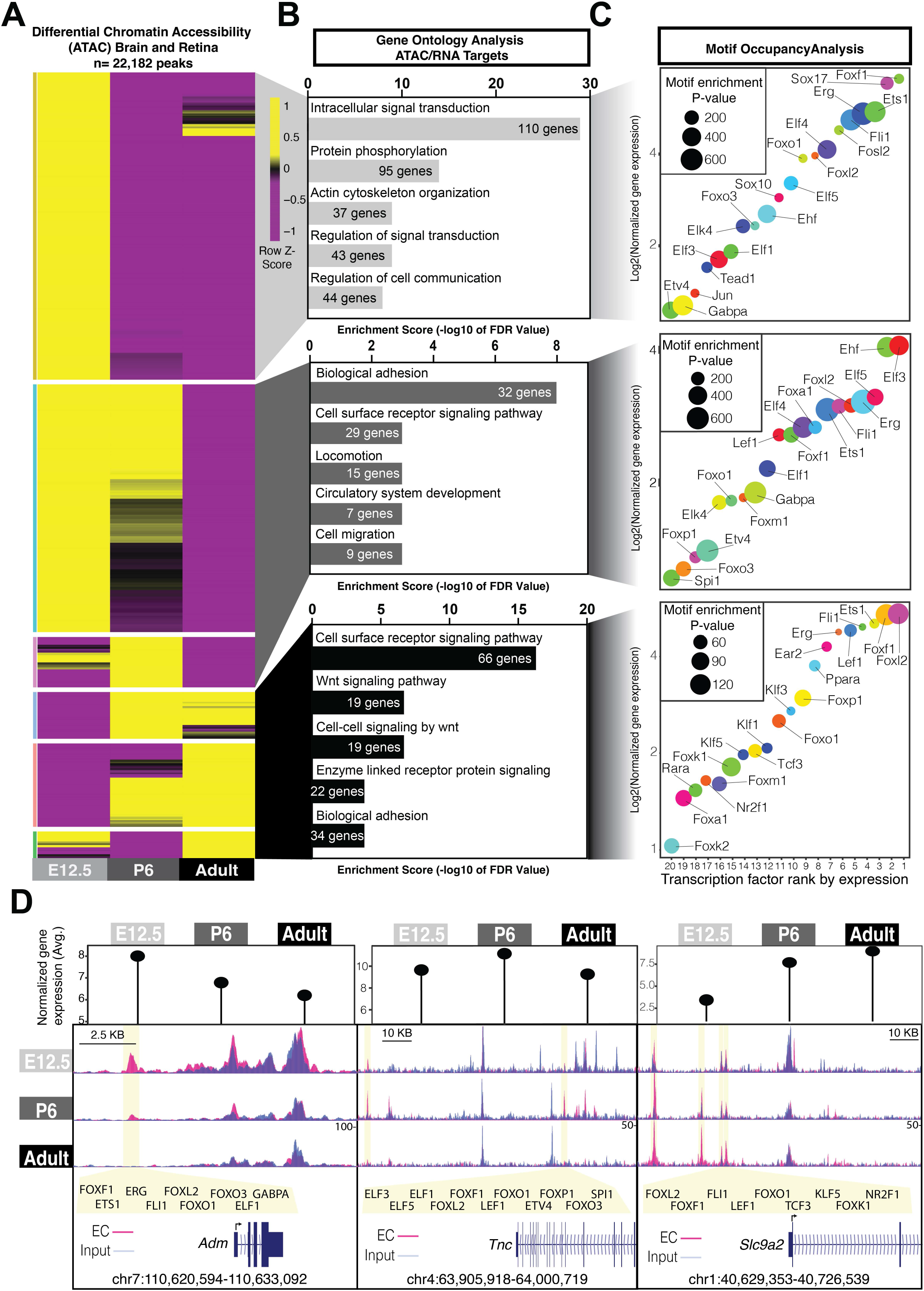
Chromatin Accessibility Changes Across Time in the Brain Endothelium. A) Differential chromatin accessibility determined by ATAC-Seq within the brain and retinal endothelium (6,540 peaks) of E12.5, postnatal day 6 (P6) and adult mice. B) Top biological processes from GREAT analysis across differentially accessible peaks at each timepoint. C) Top 20 transcription factors ranked by expression for each age. Log2 expression over input indicated in the y-axis. Motif enrichment p-value is shown according to the size of the bubble. D) Normalized gene expression in either E12.5, P6 or adult brain and retina endothelium (top) and genomic tracks for endothelial and input brain samples for genes upregulated in E12.5 (*Adm*), P6 (*Tnc*) or adult (*Slc9a2*). Unique peaks to those timepoints are indicated by the transparent vertical yellow bar, and DNA binding sites of the top 20 transcription factor motifs that are present in such peaks are indicated below.

Whereas E12.5-enriched genes showed terms related to intracellular signal transduction and actin cytoskeleton organization, postnatal day 6 (P6) endothelium was enriched for processes such as adhesion, cell surface receptor signaling, locomotion and migration. Adult-enriched CNS genes featured GO terms found at E12 and P6, such as cell surface receptor signaling pathway and biological cell adhesion, as well as novel terms related to WNT signaling and enzyme-linked receptor protein signaling (Figure 4B, Supplemental Table 4).

Next, at each timepoint we examined the most enriched transcription factor DNA-binding motifs and rank ordered them by their mRNA expression level (1=highest, 20=lowest) (Figure 4C). At E12.5, motifs for several ETS family transcription factors (ELF4, ELF5, ELK3, ELK4, etc.) were enriched in the cerebrovasculature, with ETS1, ERG, and FLI1 among the top 5 transcription factor motifs, as ranked by actual gene expression. FOXF1 and SOX17 rounded out the top 5, while other ETS, FOX and SOX family members made up the top 20, as did TEAD1 and JUN. At P6, ETS1 moved out of the top 5, and FLI1 motif enrichment was substantially decreased, while EHF, ELF3, ELF5, ERG and FOXL2 were the top 5 most enriched motifs and highly expressed transcription factors in the early postnatal CNS endothelium. In the adult CNS endothelium, FOXL2 was the most abundantly expressed of the over-represented transcription factor motifs, followed by FOXF1, ETS1, FLI1, and LEF1. LEF1 and TCF3, known regulators of canonical WNT signaling involved in blood brain barrier maturation, as well as PPARA, FOXP1, FOXO1, FOXM1, KLF1, KLF5, and NR2F1 were among the notable adult-enriched TFs (Figure 4C). Similar analysis of motif usage and transcription factor enrichment within the endothelium during development was performed for the heart (Supplemental Figure 2), liver (Supplemental Figure 3), lung (Supplemental Figure 4) and kidney (Supplemental Figure 5).

We then examined accessible, brain-specific regions of open chromatin within (or nearby) genes that were differentially expressed in E12.5, P6 or Adult CNS endothelium for these same transcription factor DNA binding motifs. *Adrenomedullin* (*Adm*), enriched in tip cells of the developing brain vasculature (Sabbagh et al., 2018), contains an accessible chromatin region in E12.5 at the zenith of *Adm* expression peaks (Figure 4D, left). Similarly, expression of *Tenascin-c* (*Tnc*), whose gene product is involved in cell adhesion (Chiquet-Ehrismann and Tucker, 2011), peaks at P6 and features two regions of open chromatin at this stage that are lost in the adult endothelium (Figure 4D, middle). Finally, *Slc9a2*, which encodes a Na/H exchanger present in brain endothelium (Lam et al., 2009), contains three regions of open chromatin upstream of its promoter that are specifically enriched in the adult endothelium (Figure 4D, right). All 3 genes contain uniquely accessible chromatin with predicted DNA binding sites for various members of the top 20 most enriched transcription factors in the brain (Figure 4D).

### Exploring Blood Brain Barrier Development at a Single Cell Resolution

Following our identification of transcription factors and their DNA binding motifs enriched in the brain endothelium by ATAC-Seq, we were interested in how these same transcriptional regulators, and their targets, changed during maturation of the CNS endothelium at a more granular level. CD31^+^ endothelial cells from whole brains (E9.5, E12.5 and E16.5), or only the cortex (P8 and Adult), were isolated by Magnetic Activated Cell Sorting (MACS) and then processed for single cell RNA-seq (scRNA-seq) (Figure 5A). After filtering (see methods), all cells isolated from E9.5 (6,039), E12.5 (6,822), E16.5 (3,358), P8 (4,048), and adult (2,723) brain were examined (Figure 5B-D, Supplemental Figure 6). As expected, dimensionality reduction and visualization of these scRNA-seq data by uniform manifold approximation and projection (UMAP) revealed a fairly uniform distribution of cells between samples (Hao et al., 2021; Melville, 2020) (Figure 5B). Cell identities were assigned based on the expression of well-characterized marker genes, with astrocyte, microglia, mural, and macrophage populations identified within our brain datasets (Figure 5C, Supplemental Figure 6B-D, Supplemental Table 5). Endothelial cell clusters, expressing characteristic EC transcripts such as *Cdh5*, were evident at all stages examined, validating the CD31^+^ MACS enrichment (∼79% of the 22,990 sequenced cells were endothelial cells) (Figure 5D).

**Figure 5:**
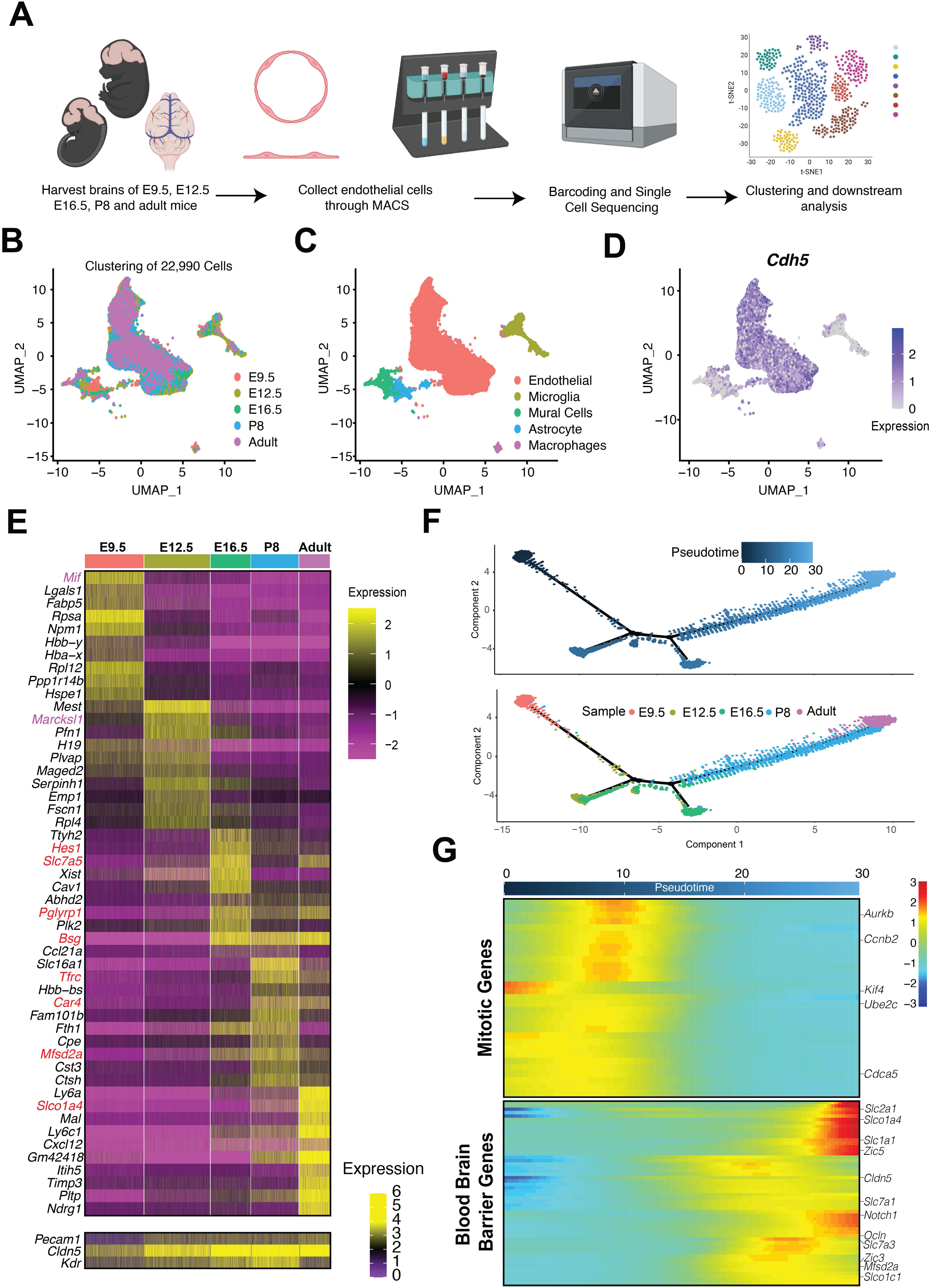
Maturation of Blood Brain Barrier at Single Cell Resolution. A) Schematic representation of the harvesting and isolation of endothelial cells from E9.5, E12.5, E16.5, P8 and adult mice. Cells were purified using Magnetic Isolation Cells Sorting (MACS) and processed for downstream sequencing and analysis following the 10x Genomics protocol. B) UMAP representation of total cells sequenced from all timepoints. C) Clustering annotation and identity of the cell types sequenced. D) Feature plot showing *Cdh5* expression enrichment in the endothelial cell cluster. E) Heatmap of differential gene expression analysis of endothelial cells from each timepoint. Genes in red have a known role in blood brain barrier function. Top 10 genes are shown, followed by *Pecam1*, *Cldn5* and *Kdr*. F) Monocle pseudotime analysis of all endothelial cells from all timepoints, with E9.5 set as the point of origin. The pseudotime gradient is shown on top and the corresponding timepoints are color coded below. G) Heatmap showing expression dynamics of selected gene markers for mitosis or blood brain barrier development markers superimposed on the pseudotime axis.

To define gene expression changes within brain endothelial cells over time, the endothelial cluster was extracted and further analyzed. Differential gene expression signatures were evident between the various time points (Figure 5E, Supplemental Table 5). *Macrophage migration inhibitory factor* (*Mif*), an inflammatory cytokine with chemokine functions that has been implicated in angiogenesis (Amin et al., 2003), was robustly expressed in E9.5 brain endothelial cells, but markedly downregulated in later stages. *Marcksl1*, a gene involved in blood vessel shape and size (Kondrychyn et al., 2020), was the most differentially upregulated gene in the E12.5 brain endothelium (*Mif* and *Marcksl1* are labeled in purple, Figure 5E), while the amino acid transporter *solute carrier transporter 7a5* (*Slc7a5*) (Tarlungeanu et al., 2016), as well as other blood brain barrier markers (denoted in red), initiated expression at E16.5 when blood brain barrier formation begins (Ben-Zvi et al., 2014; Hupe et al., 2017). Conversely, expression of *plasmalemma vesicle-associated protein (Pvlap/ Mecca 32)*, a pan-endothelial marker that is lost in the mature BBB endothelium (Benz et al., 2019; Guo et al., 2016), was dramatically decreased after E12.5. *Major facilitator super family domain containing 2a (Mfsd2a)*, which encodes a lipid transporter required for proper blood-brain barrier development (Ben-Zvi et al., 2014; Wong and Silver, 2020), and *solute carrier organic anion transporter family member 1a4* (*Slco1a4*), an organic anion transported recently studied as a potential target for drug delivery to the brain (Akanuma et al., 2013; Ose et al., 2010), are both enriched E16.5 through adult brain endothelium.

Next, we performed pseudotemporal ordering of individual CNS ECs to further characterize their developmental trajectories (Qiu et al., 2017a; Qiu et al., 2017b; Trapnell et al., 2014) (Figure 5F). Genes involved in mitosis, cell division and proliferation, such as *Aurora Kinase B* (*Aurkb*) (Bischoff and Plowman, 1999; Giet and Prigent, 1999), *Kinesin superfamily protein 4* (*Kif4*) (Hu et al., 2011), and *Marker of proliferation Ki-67* (*Mki67*) (Booth et al., 2014), were markedly elevated in early brain development, when angiogenesis is rapidly expanding the vascular network. Conversely, at the other end of the pseudo time spectrum, genes involved in blood brain barrier maturation, such as the tight junction encoding genes *Claudin 5* (*Cldn5*) (Nitta et al., 2003) and *Occludin* (*Ocln*) (Argaw et al., 2009), as well as the transporters *Mfsd2a* (Ben-Zvi et al., 2014; Wong and Silver, 2020) and *Glut1* (*Slc2a1*) (Veys et al., 2020) initiated at E16.5 and peaked in the P8 and adult endothelium (Figure 5G).

### Identification of Gene Regulatory Networks Involved in Brain Endothelial Development

To identity potential transcriptional regulators of cerebrovascular development and maturation we utilized Single-Cell rEgulatory Network Inference and Clustering (SCENIC) (Aibar et al., 2017). By correlating transcription factor expression within individual endothelial cells along with expression of their presumptive targets, SCENIC predicts active gene regulatory networks (GRNs). First, sets of genes that are co-expressed with transcription factors are identified as a module. Then, putative direct-binding targets within a module are examined for the presence of cis-regulatory motifs of these co-expressed transcription factors to generate a “regulon”, while indirect targets are removed. This process is repeated for each transcription factor, and its putative co-expressed targets, expressed within each cell. Finally, cells with similarly active regulons (or GRNs), are then grouped together (Figure 6A).

**Figure 6:**
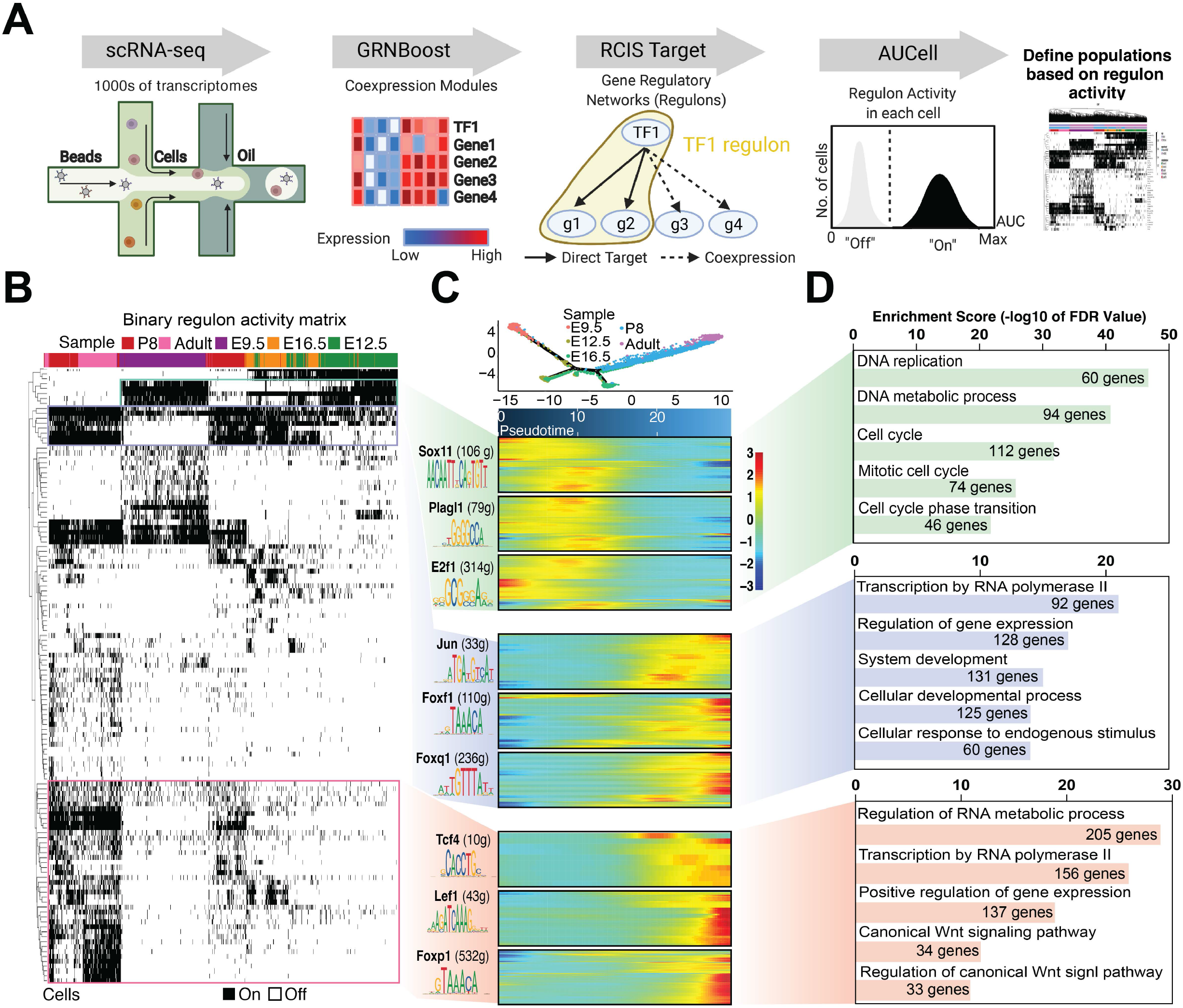
Gene Regulatory Networks Involved in Blood Brain Barrier Development. A) The SCENIC (Aibar et al., 2017) analysis pipeline. scRNA-Seq co-expression modules between (1) TFs and (2) candidate target genes are inferred using GRNBoost. RCis Target then identifies modules for transcription factor DNA-binding motifs that are enriched across the target genes to create a “regulon” of direct targets. AUCell scores the activity of each regulon in every single cell, generating a binary activity matrix to predict cell states. B) SCENIC binary activity heatmap representing active regulons in brain endothelial cells across all timepoints. Vertical columns represent individual sequenced cells, while each horizontal row represents an individual regulon. Highlighted regulons are shown in panel C. C) Heatmaps show differentially active regulon target gene expression in the cerebral endothelium at E9.5 and E12.5 (green shading) compared to E16.5, P8 and adult (blue shading) and P8 and adult (orange shading), all superimposed upon the pseudotime gradient from Figure 6G. D) Selected GO biological processes derived from the target genes expressed in each of the three regulon clusters shown in panel C.

Using SCENIC, we identified 3 distinct endothelial clusters based upon regulon activity (Figure 6B). The first cluster of regulons, including SOX11 (106 genes), PLAGL1 (79 genes) and E2F1 (314 genes), are enriched primarily in the E9.5 and E12.5 brain endothelium. SOX11 regulates vascular development and is active during pathological angiogenesis (Palomero et al., 2014; Schmitt et al., 2013), while PLAGL1 controls early developmental angiogenesis (Starks et al., 2020), and E2F1 modulates vascular endothelial growth factor (VEGF) expression (Qin et al., 2006). Visualizing the direct transcriptional targets of SOX11, PLAGL1, and E2F1 in context of CNS EC brain maturation using pseudotime analysis revealed that these putative gene regulatory networks were largely upregulated in immature endothelia (e.g. E9.5), while GO analysis showed their target genes are involved in DNA replication and the cell cycle (Figure 6C, D). The second cluster of regulons identified by SCENIC were active primarily in the E16.5, P8 and adult CNS endothelium, including JUN (33 genes), FOXF1 (110 genes) and FOXQ1 (236 genes). *Jun* has been implicated in tip cell specification and tube formation during angiogenesis (Keisuke et al., 2020; Licht et al., 2006; Yoshitomi et al., 2021). *Foxf1* is critical for endothelial barrier function in the lung, but is not required for blood brain barrier maintenance (Cai et al., 2016), while *Foxq1* is enriched in the developing brain endothelium (Hupe et al., 2017). Gene ontology predicts that transcripts in this second cluster are involved in processes such as the regulation of gene expression, system development and cell proliferation. The third and last cluster identified by SCENIC contained regulons active in the P8 and the adult CNS endothelium, including TCF4 (10 genes), LEF1 (43 genes) and FOXP1 (532 genes). *Lef1*, which encodes an obligate binding partner of β-catenin in the nucleus, as well as *Tcf4* (*Transcription factor 4*) both act downstream of canonical WNT signaling to govern blood brain barrier function (Wang et al., 2019; Zhou et al., 2014). GO analysis shows target genes downstream of these adult enriched transcription factors were involved in macromolecule modification, regulation of cellular metabolic processes, and WNT signaling (Figure 6C-D). Furthermore, some target genes were present in more than one regulon, suggesting they may function as critical nodes in brain endothelial development (Supplemental Figure 7, full list in Supplemental Table 6). Notably, many of the GRNs identified by SCENIC featured enriched DNA binding motifs and upregulated gene expression for transcription factors identified in our ATAC-seq and RNA-seq analysis, such as JUN, FOXF1, and LEF1 (Figure 4C). Interestingly, *Nuclear Receptor Subfamily 3 group C member 1* (*Nr3c1*), which encodes a glucocorticoid receptor and is involved in the regulation of WNT/β-catenin pathway (Liu et al., 2021) and *albumin D-binding protein (Dbp)*, a proline amino-acid-rich domain basic leucine zipper (PAR bZip) transcription factor involved in circadian rhythm control in the blood brain barrier (Franken et al., 2000; Pulido et al., 2020), also showed an increase in regulon activity (Supplemental Figure 8A-C).

### Cell Type Specific Regulon Activity in the Cerebrovasculature

An advantage of scRNA-Seq is that it enables the identification of distinct endothelial cell types based on marker gene expression, allowing one to distinguish between various endothelial identities, such as arterial, capillary-arterial, capillary-venous, venous, mitotic and tip cells (Sabbagh et al., 2018; Vanlandewijck et al., 2018). Given the dynamic nature of LEF1 and FOXP1 regulon activity within the brain vasculature during development, we wondered if these gene regulatory networks were uniformly active across all vessel types (Figure 7A). To detect changes in regulon activity at different developmental timepoints, we first subclustered E12.5 and adult brain ECs using defined markers for these different vessel identities (e.g. arterial, capillary vein, capillary artery, venous, tip cell, and mitotic) (Sabbagh et al., 2018). Both E12.5 and adult CNS ECs contained cells from each unique vessel identity (Figure 7B-E). Interestingly, whereas the LEF1 regulon was enriched in tip cells and capillaries at E12.5, it expanded to encompass all vessel types in the adult brain (Figure 7F). Conversely, the FOXP1 regulon was selectively active in arterial cells at E12.5, but it also expanded to include all vessel types in the adult brain (Figure 7G).

**Figure 7:**
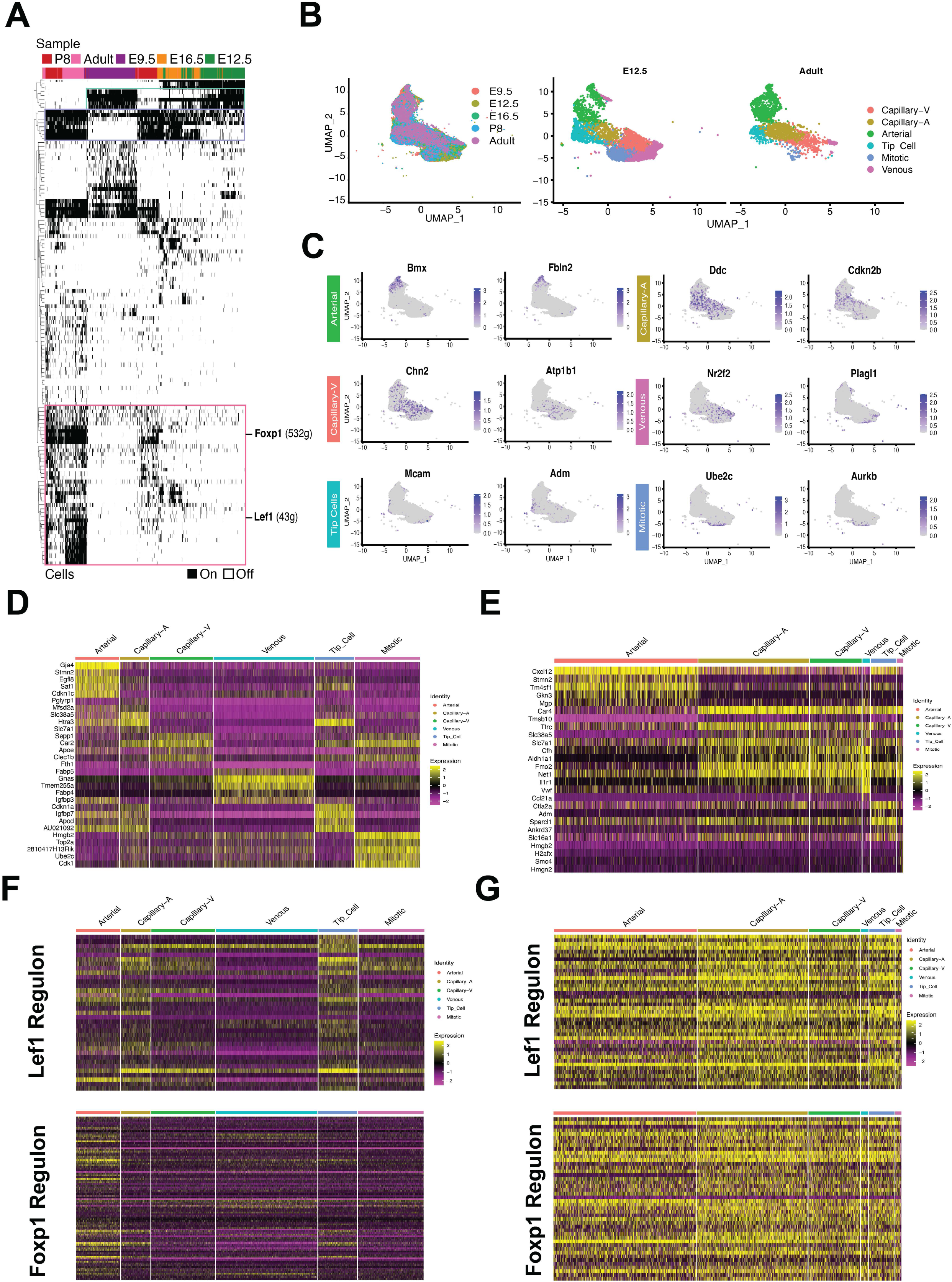
Vessel Specific Changes in Regulon Activity in the Brain Endothelium During Development. A) SCENIC binary activity heatmap representing active regulons across endothelial cell timepoints, with the FOXP1 and LEF1 regulons active in the P8 and adult brain endothelium indicated on the right. B) UMAP representation of all endothelial cells labelled by timepoint (left) and by endothelial subtype corresponding to arterial ECs, capillary-arterial (Capillary-A), capillary-venous (Capillary-V), venous, mitotic and tip-cells at E12.5 (middle) and in the adult brain (right). C) Feature plot showing expression of marker genes with enriched expression in each cluster. A heatmap shows the top 5 differentially expressed genes from each cluster from the E12.5 (D) and adult (E) brain endothelium. Heatmaps showing expression of LEF1 and FOXP1 regulon targets in the E12.5 (F) and adult (G) brain endothelium.

### Neurovascular Unit Interactions Change During Blood Brain Barrier Maturation

The blood brain barrier is part of the neurovascular unit (NVU), which is composed of neurons, mural cells (i.e. smooth muscle, pericytes), glia and astrocytes that surround and interface with the cerebral endothelium (Schaeffer and Iadecola, 2021). Using NicheNET (Browaeys et al., 2020), we next identified ligands expressed in non-EC cells of the NVU within our dataset, as well as their target genes expressed in the CNS endothelium, to determine whether these ligand-target interactions are driving activation of the regulons identified by SCENIC within the brain vasculature. We examined only genes that were significantly upregulated in the adult endothelium compared to the embryonic day 9.5 (E9.5) endothelium, and with endothelial cells designated as the signal receiving cells (receptors and downstream effectors), with other cell types of the NVU (microglia, pericytes and mural cells) being defined as signal sending cells. From this analysis we identified the upregulation of cell adhesion molecules in the endothelium, such as *catenin delta 1* (*Ctnnd1, P120)* (Anney et al., 2021) (Supplemental Figure 9A). Expression of *Ctnnd1*, along with WNT signaling regulated genes such as *Cyclin dependent kinase inhibitor 1A* (*Cdkn1a*) (Nayak et al., 2018), *Cyclin D1* (*Ccnd1*) (Shtutman et al., 1999; Tetsu and McCormick, 1999), *Prothymosin Alpha* (*Ptma*) (Lin and Chao, 2015), and *Catenin beta-1/ β-catenin* (*Ctnnb1*) were predicted to be induced by pericyte-mediated presentation of the ligand Cadherin 2 (CDH2) to the endothelium (Ortiz et al., 2015; Zheng et al., 2016). Furthermore, pericyte CDH2 can also induce endothelial expression of *Lef1* (Soh et al., 2014) and the canonical Wnt target, *Axin2 (Jho et al., 2002).* Importantly, endothelial expression of VE-Cadherin (*Cdh5)* can also induce *Lef1* (Birdsey et al., 2015). Genes involved in vascular maintenance, such as *Rad51*, are potentially driven by SMC expression of *Integrin beta 1* (*Itgb1*) (Ahmed et al., 2018; Vattulainen-Collanus et al., 2018) (Supplemental Figure 9A).

After identifying the putative downstream effectors within endothelial cells induced by ligands expressed in neighboring cell types of the NVU, we next focused on the ligands presented by the endothelium and their potential receptors in pericytes, which stabilize capillary vessels in the brain (Supplemental Figure 9B). Using CCInx (version 0.5), we found that the adult cerebral endothelium is enriched for chemokines that regulate leukocyte migration and maintain homeostasis, such as *Cxcl12*, while its receptor, *Ackr3/Cxcr7*, is enriched in pericytes (Boldajipour et al., 2008; Williams et al., 2014). Similarly, adult brain ECs express *Pdgfb*, while its cognate receptor, *Pdgfrb*, was enriched in adult pericytes (Abramsson et al., 2007; Gaengel et al., 2009). An EC to pericyte interaction was also noted for *Amyloid precursor protein* (App) and *Vitronectin* (*Vtn*) (Calero et al., 2012). Conversely, the adult brain endothelium featured decreased expression of *Macrophage migration inhibitory factor* (*Mif*), which is known to reduce pericyte contractility (Pellowe et al., 2019), while pericytes decreased expression of multiple potential MIF receptors, including *Transferrin Receptor 1* (*Tfrc*, *Cd71*) and *Integrin α4* (*Itgα4*). Collectively, these data show cellular communication within the NVU can be readily inferred from scRNA-seq data within the developing murine brain, as both known and novel interactions were evident between ECs and mural cells.

### Identification of Evolutionarily Conserved Regions of Open Chromatin

To investigate if the transcription factor networks we identified in the murine brain play an analogous, conserved role in humans, we turned to an *in vitro* model of the human brain vasculature: hCMEC/D3 cells (Weksler et al., 2013). Using Omni-ATAC-seq (Corces et al., 2017), regions of open chromatin were identified in these cultured human brain endothelial cells and then compared to accessible regions within the P8 and adult murine brain. Of the 94,197 regions of open chromatin identified in human brain microvascular endothelial cells, 15,131 were conserved in the mouse genome (mm10). Out of these evolutionarily conserved regions, 314 overlapped with regions that were uniquely accessible within the adult murine brain endothelium (Figure 8A, Supplemental Table 7), and the most enriched transcription factor DNA binding motifs within these conserved, accessible regions was determined using HOMER (Figure 8B). Notably, common core endothelial TF motifs, such as ETS DNA binding sites, did not emerge at the top of this list as this analysis focused on regions and motifs that were enriched specifically within the endothelium of the postnatal and adult brain. Transcription factor motifs that were evolutionarily conserved in the open chromatin of the adult human and murine cerebral endothelium were FOXM1, FOXL2, FOXA1, FOXF1, and BATF. Interestingly, conserved regions of open chromatin that mapped to genes expressed in both human and murine brain vasculature (via GREAT and RNA-Seq) were involved in processes such as vascular development, cell communication, and WNT signaling (Figure 8C). Examples of these evolutionarily conserved, putative regulatory elements in the adult cerebral endothelium can be found within the first intron of *Slc31a1* (*Solute Carrier Family 31 Member 1*), which contains motifs for TCF4, LEF1 and FOXO3, and in a region proximal to *Mfsd2a* (*Major facilitator superfamily domain-containing protein 2*), that has motifs for TCF4, LEF1 and ETS (Figure 8D). *Msfd2a* encodes for a critical lipid transporter that is enriched in the brain endothelium (Andreone et al., 2017; Ben-Zvi et al., 2014; Nguyen et al., 2014; O’Brown et al., 2019), and loss of WNT signaling either in receptor (*Lrp5^-/-^*) or ligand (*Ndp^y/-^*) mice downregulates *Mfsd2a* expression and increases transcytosis and BBB breakdown in mice (Wang et al., 2020).

**Figure 8:**
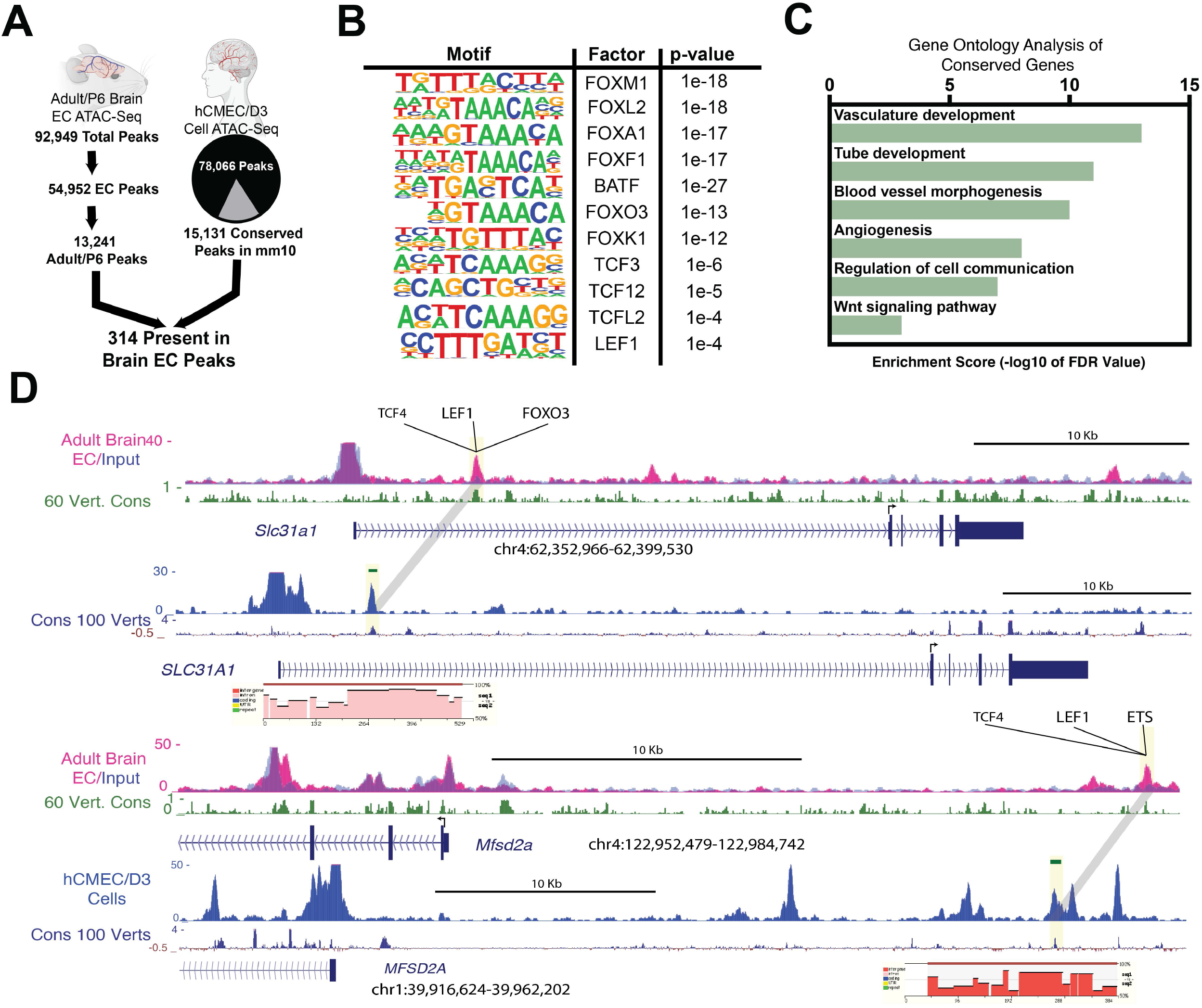
Evolutionary Conservation of Regions of Open Chromatin Between Human and Adult Mouse. A) Diagram representing the total number of open chromatin regions in hCMEC/D3 that are conserved in the adult murine brain endothelium (shown in Figure 4). B) Selected known transcription factor DNA binding motifs in conserved peaks along with their p-value after analysis by HOMER. C) GO term analysis of genes with conserved nearby accessible chromatin regions that are also expressed in both human hCMEC/D3 cells and adult murine brain endothelium. D) Representative genome browser tracks of *Slc31a* and *Mfsd2a* highlighting (in yellow) conserved accessible chromatin regions in human (top) and murine (bottom) as defined by ATAC-seq and Omni-ATAC-seq. Transcription factor motifs present in the highlighted peak are shown above. Conservation at the nucleotide level within each highlighted peak is shown below each locus.

## DISCUSSION

Herein, we have profiled the accessible chromatin and gene expression signatures of the embryonic, postnatal, and adult brain and heart, as well as the postnatal and adult retina, liver, kidney, and lung endothelium. By establishing a lexicon of common, accessible regions of open chromatin present within the endothelium of these six organs, across developmental time, we have identified a core set of enriched transcription factor DNA binding motifs common to all endothelial cells, regardless of their origin. Additionally, we extend these observations to identify accessible regions in the genome that are enriched in specific organs, along with the possible transcription factors that act on these putative regulatory elements to give rise to the functional heterogeneity evident within these different vascular beds (Sabbagh et al., 2018). Moreover, using single cell transcriptomic approaches we interrogate the gene regulatory networks governing development and maturation of the cerebrovasculature at the single cell level. Finally, we demonstrate that the regulatory regions, and the transcription factor motifs within them that we identified in the adult murine CNS endothelium are evolutionary conserved in humans.

Significantly, within these accessible regions of open chromatin within the endothelium, the DNA binding motif for the ETS family of transcription factors are the most commonly occurring TF binding site, regardless of organ identity. This was expected, given the key functions ETS TFs play in endothelial specification, vessel growth, and angiogenesis (Asano et al., 2010; Birdsey et al., 2015; De Val and Black, 2009; Palikuqi et al., 2020). Other common, core motifs present in the endothelium of all organs were those of the SOX transcription factor family (Chiang et al., 2017; Yao et al., 2019b). Critically, organ-specific signatures also emerged, as analysis of open chromatin unique to the vasculature of each organ identified an array of transcription factor binding motifs enriched to each tissue, such as GATA4 in the liver, and NFAT in the heart. While we focused our attention on the cerebrovasculature, this catalogue of chromatin landscapes and gene expression signatures of the endothelium of different organs is a valuable resource that can be further interrogated to generate new hypotheses regarding endothelial specialization, maturation, and homeostasis.

The mature brain vasculature features unique characteristics, such as extensive cell-cell junctions, and selective permeability (Obermeier et al., 2013). This specialization, along with the need to define the transcriptional networks governing the establishment and maintenance of the blood brain barrier, warranted further investigation at the single cell level over developmental time. Examination of 18,827 single bran endothelial cell transcriptomes, across 5 distinct developmental stages, revealed a stark transition from a mitotic, and proliferative signature at E9.5, to a homeostatic endothelium featuring a rich repertoire of channels and transporters evident in the adult brain. This was expected, as the predominant mechanism of early blood vessel growth within the brain is angiogenesis (proliferation, migration, sprouting), while growth begins to wane as the existing capillaries and larger diameter vessels mature and remodel to establish the blood brain barrier from E16.5 through postnatal development. Critically, using scRNA-seq we identified novel GRNs in the early brain, such as SOX11, PLAG1, and E2F1, while also showing confirming our ATAC-seq and RNA-seq results which suggested that JUN, FOXF1, and FOXQ1 control maturation of the brain endothelium. Critically, AP-1 transcription factors, such as JUNB, control vascular development in the retina (Engelbrecht et al., 2020; Keisuke et al., 2020). Whether other TFs and their GRNs identified herein, such as FOXF1, interact with the WNT signaling pathway to regulate BBB maturation remains unknown (Ustiyan et al., 2018). Finally, our single cell data also identified robust LEF1, NR3C1, and DBP regulons specific to the adult brain endothelium. Identification of a LEF1 GRN within the adult brain vasculature consistent with recent studies demonstrating a critical requirement for *Lef1* in blood brain barrier maturation (Daneman et al., 2009; Mike et al., 2017; Roudnicky et al., 2020; Zhou et al., 2014). However, our temporal and cell type specific analysis revealed that a LEF1 GRN is, in fact, active in early tip and capillary cells of the early cerebral endothelium, and it then expands during development to become upregulated in all vessel types within the postnatal brain. A similar pattern, albeit being confined to the early arterial endothelium, was evident for the FOXP1 GRN. While there are fewer links in the current literature between either DBP or NR3C1 and the CNS vasculature, reports do suggest *Dbp* and its transcriptional targets control circadian rhythms within the CNS (Lopez-Molina et al., 1997; Pulido et al., 2020), and some studies suggest NR3C1 plays a role in vascular inflammation and aneurysm (Al Argan et al., 2018; Goodwin et al., 2015). Of interest will be future studies of these same GRNs in neurovascular diseases accompanied with BBB disruption.

Finally, by performing a cross-species analysis to another vertebrate, our data demonstrate the major DNA binding motifs found in the murine adult cerebrovasculature were also present within a human cell culture model of the blood brain barrier. Similar to what was observed in our murine dataset, the genes linked to these evolutionarily conserved, accessible chromatin regions in the human brain endothelium were also involved in blood vessel morphogenesis and WNT signaling. These conserved regions are of great interest, and future studies will interrogate the sufficiency and necessity of these potential brain specific enhancers to modulate gene expression *in vivo*.

### Limitations of the Present Study

Changes in open chromatin do not directly translate to changes in gene expression. Furthermore, the chromatin surrounding most proximal promoters are likely in an accessible state in most situations, as the transcriptional status of many loci is not determined by differential accessibility, per se, but by differential recruitment of the transcriptional machinery, or even post-translation modification of already engaged protein complexes (as occurs in pause-release of the Pol II transcriptional machinery at the proximal promoter) (Adelman and Lis, 2012; Fish et al., 2017; Jonkers and Lis, 2015; Narita et al., 2021). A technical limitation of our work is the methods and analysis used herein infer enhancers of target genes, rather than measure direct looping or physical contacts (e.g. as in chromatin conformation capture techniques). Moreover, these putative enhancers, as well as the novel gene regulatory networks identified by scRNA-seq, have not been functionally validated. Critically, bulk nuclear RNA-Seq yielded less robust transcript number than traditional bulk whole cell RNA-Seq. Whether this was due to loss of cytoplasmic RNA, or inadequate input material, is unknown. Finally, our *in vitro* chromatin accessibility data from cultured human microvascular endothelial cells likely does not fully reflect the transcriptional complexity of the intact adult human brain.

## Conclusion

In summary, we present a comprehensive catalogue of the chromatin landscape within the endothelium of multiple organs of the developing and adult mouse. This data is augmented by a granular dissection of the development and maturation of the brain endothelium, and the gene regulatory networks acting at the level of single cells within this organ. Finally, we demonstrate that many of these accessible regions of open chromatin, and the DNA binding motifs contained within these regions, are well conserved between mice and humans. By studying the unique chromatin landscape of healthy endothelial cells throughout the organs of the body, this resource will guide future studies aimed at experimentally manipulating these unique populations, and it suggests novel targets for promoting engraftment of new endothelium within each organ.

## MATERIAL AND METHODS

### Mice

All mouse protocols were approved by the Institutional Animal Care and Use Committee (IACUC) at Baylor College of Medicine. For all experiments, noon on the day a vaginal plug was discovered was considered embryonic day 0.5, the day of birth was considered P0, and all adult mice were 8 weeks of age.

### Genotyping and mice used

*Cdh5(PAC)-CreERT2* mice (MGI #: 3848982) were from Ralf Adams. *Rosa26-Sun1-sfGFP-6xMyc* (e.g. *R26^Sun1GFP^)* (MGI #: 5443817) were purchased from Jax. Genotyping for all alleles was performed by PCR using gene specific primers. Please see Supplemental Materials and Methods for more details.

### Murine Endothelial Nuclear isolations

For embryonic analysis, tamoxifen (0.015 mg/kg bodyweight) was administered to pregnant dams by intraperitoneal (i.p.) injection at E10.5 and embryos were collected at E12.5. For postnatal tissues, tamoxifen (0.015 mg/kg bodyweight) was administered by subcutaneous injection at P1 and P3, and tissues were collected at P7. For adult experiments, tamoxifen (0.015 mg/kg bodyweight) was administered by i.p. injection 7 days prior to tissue isolation. In all cases, after gross dissection, GFP expression within the vasculature of each tissue of interest (or embryo) was confirmed by direct immunofluorescence for each sample collected. GFP negative samples were not processed further. Nuclear isolation was performed according to Mo et. al (Mo et al., 2015). Briefly, fresh tissue was harvested on ice in Buffer HB++ composed of 0.25 M sucrose, 25 mM KCl, 5 mM MgCl_2,_ 20 mM Tricine-KOH, pH 7.8 with protease inhibitors (Roche/Sigma Cat. #11873580001), 1 mM DTT (Sigma D0632), 0.15 mM Spermine (Sigma S1141), 0.5 mM Spermidine (Sigma S2501), and RNAse inhibitors (Promega N2611) and immediately dissected and minced into 1 mm-by-1 mm portions with curved scissors. Tissue was transferred along 1ml of HB++ in a chilled Eppendorf tube in ice and homogenized using Bio-gen Series PRO200 homogenizer. Short bursts of ∼5-8 seconds were done to prevent overheating. Once no large pieces were observed, the tissue was transferred to large clearance dounce homogenizer “A” (7ml, Kontes Glass Company) and 4 mL of HB++ was added. Tissue was homogenized with 20 strokes and transferred to small clearance homogenizer “B”, 320 ul of 5% IGEPAL CA-630++ in HB++ was added and dounced with the tight pestle 20 more times slowly to avoid creating bubbles and disrupting cell membranes. The homogenate was then strained using a 40 μm cell strainer into a 50 mL conical tube. 5 mL of working solution of 5 volumes of Optiprep solution (Sigma, D1556) and one volume of diluent (150 mM KCl, 30 mM MgCl_2,_ 120 mM Tricine-KOH, pH 7.8 in water) was added and homogenized by inversion and poured into an empty pre-chilled 30 mL Corex tube. Once all samples were ready, using a pipette aid, the tip was placed just above the bottom surface of the Corex tube, and sample was slowly underlying with 7.5 mL of the 30% and then 4 mL of the 40% iodixanol++ solutions (diluted with buffer HB). Nuclei were then isolated by density gradient centrifugation with optiprep density gradient medium. Nuclei were collected from the 30-40% interface and then pre-cleared with Protein-G Dynabeads (Life technologies, 10003D). A portion of these nuclei were held back for use as input samples. Next, GFP^+^ nuclei were immunoprecipitated with an anti-EGFP antibody (ABfinity Rabbit monoclonal anti-GFP antibody; 0.2 mg/mL) for 40 minutes at 4°C with gentle agitation, followed by binding to Protein-G Dynabeads (Invitrogen, 10003D) for 20 minutes hours at 4°C to enrich for endothelial cell nuclei. Isolated nuclei were filtered using 20 μm Celltrics filter (Sysmex #04-004-2326).

Specific amounts of tissue and yields of nuclei from each tissue are listed below. For adult hearts, 4 hearts were used per INTACT experiment with 80% of the tissue processed resulting in a total of 1.07×10^6^ isolated nuclei. For adult lungs, 2 lungs per INTACT experiment were used with 60% of the tissue processed and resulting in a total of 1.1×10^6^ isolated nuclei. 10 adult retinas were used per INTACT experiment resulting in 50,000 isolated nuclei. 1 adult brain was used per INTACT experiment with resulting in 1.45 x10^6^ isolated nuclei. 1 adult liver was used per INTACT experiment with 50% of the tissue processed resulting in 5 x10^5^ isolated nuclei. 4 adult kidneys were used per INTACT experiment with 60% of the tissue processed and resulting in 8.5×10^5^ isolated nuclei. 8 P7 hearts were used per INTACT experiment resulting in 9×10^5^ isolated nuclei. 4 P7 lungs were used per INTACT experiment resulting in 395,000 isolated nuclei. 16 P7 retinas were used per INTACT experiment resulting in 85,000 isolated nuclei. 6 P7 brains were used per INTACT experiment with 60% of the tissue processed and resulting in 1×10^6^ isolated nuclei. 8, P7 livers were used per INTACT experiment with 60% of the tissue processed and 6.7×10^5^ isolated nuclei. 8, P7 Kidneys were used per INTACT experiment resulting in 1.2×10^6^ isolated nuclei. 5 E12.5 trunks per INTACT experiment were used per INTACT experiment resulting in 1.0 x10^6^ isolated nuclei. 5 E12.5 brains were used per INTACT experiment resulting in 1.5 x10^5^ isolated nuclei. 5 E12.5 hearts were used per INTACT experiment resulting in 47,000 isolated nuclei. Each isolation was performed at least twice.

### Assay for Transposase-Accessible Chromatin with high throughput sequencing (ATAC-seq)

Approximately 50,000 bead-bound EGFP^+^ and 50,000 input nuclei from murine tissues were used as input for ATAC-seq. ATAC-seq libraries for murine endothelial cells were processed as previously described (Buenrostro et al., 2015) and libraries were generated using the Nextera DNA Sample Preparation Kit (Illumina, FC-121-1030). The quality of purified DNA libraries was checked by Agilent High Sensitivity DNA kit (Agilent Technologies). Paired-end, 2 x 75 bp sequencing was performed on an Illumina Nextseq 500 instrument. Reads were mapped to the mm10 version of the mouse genome using Bowtie2 with default paired-end settings (Langmead and Salzberg, 2012). Mitochondrial reads, reads with a MAPQ < 10, and reads which did not align to the reference genome were removed using Samtools (version 1.13) (Danecek et al., 2021). Duplicated reads were then removed with Picard MarkDuplicates (Institute, 2019). Peak calling was carried out with MACS2 (callpeak --nomodel –broad) (v2.2.7.1)(Zhang et al., 2008). Diffbind (version 3.2) (Ross-Innes et al., 2012; Stark R, 2011) was used to import peaksets (min.overlap= 0.66) into RStudio Server (version 1.4.1717, https://www.rstudio.com) using R (version 4.1, (Team, 4.1). The dba.blacklist function was used to filter out peaks that overlap with the ENCODE blacklist. Differentially accessible regions between the endothelium and the input nuclei of each organ were extracted using DESeq2 (version 1.32.0) (Love et al., 2014) with <p-value 0.5 and >1 fold change difference. Endothelial-enriched peaks from each organ were compared using the mergepeaks function in Homer (version 4.11) (Heinz et al., 2010). Peaks present in all organs were used for analysis in Figure 2. Peaks present in single organs were used for analysis in Figure 3 and Supplemental Figures 3-6. Motif enrichment analysis was conducted with findMotifsGenome and enrichment graphed as previously described (Liu et al., 2019). Graphs for individual motif distance from peaks were created using annotatePeaks in Homer and presented in an enrichment plot (Liu et al., 2019). Gene ontology analysis was done using GREAT (version 4.0.4) (McLean et al., 2010).

### Nuclear RNA-seq

In parallel to our ATAC-seq experiments, all remaining bead-bound EGFP^+^ nuclei were processed for RNA extraction using the RNeasy Plus Micro kit (Qiagen). Nuclear RNA-seq libraries were constructed with the Stranded RNA-seq Kit with Ribo Erase (Kapa Biosystems, KK8484) with custom Y-shaped adapters. Paired-end 2 x 75 bp NSQ 500/550 Hi Output KT v2.5 −75 CYS (Illumina, 20024906) was performed for RNA-seq libraries on an Illumina Nextseq 500 instrument. Reads were first mapped to the mouse genome (mm10) using Salmon (version 1.5.1) (Patro et al., 2017). Transcript level quantification was then imported using txtimport (version 1.20.0) (Soneson et al., 2015) and analyzed using DESeq2 (Love et al., 2014). Differentially expressed genes between the endothelial and input nuclei were defined as those transcripts with an expression log_2_fold-change >0.5 and Benjamini-Hochberg adjusted p-value (q-value) < 0.1. Volcano plots were created using EnhancedVolcano (version 1.10.0) (Blighe K, 2021).

### Magnetic Activated Cell Sorting for Murine Single Cell RNA-Sequencing

Brain tissue was processed for CD31 MACS with slight variations depending on the time point analyzed. For embryonic brains (E9.5, E12.5, E16.5), embryos were harvested in ice-cold Buffer HBSS++ (HBSS plus FBS, pen/strep, and HEPES). Dissected brains were placed in 250 μL of Collagenase (1 mg/mL) and placed at 37°C for 15 minutes. Tissue was pipetted up and down every two minutes, first with a P1000, then with a P200, until few to no clumps of tissue were visible. For P8 and adult brain, the cortex was dissected, and cells were dissociated using the neural tissue dissociation kit P (Miltenyi, 130-092-628). For all time points, the cell suspension was pelleted (5 min, 800 x g), then washed two times with PBS, and then resuspended in 180 μL MACS PEB buffer (phosphate-buffered saline (PBS), pH 7.2, 0.5% bovine serum albumin (BSA), and 2 mM EDTA. The cell suspension was then incubated for 15 minutes with 20 μL of anti-CD31 MicroBeads (Miltenyibiotec, Cat. No. 130-097-418) at 4°C. Cells were then washed with 1 mL of PEB buffer, centrifuged at 300 x g for 5 minutes, and applied to an MS Column (Miltenyi, 130-042-201) on a magnetic stand. After three consecutive washes on a magnetic stand with PEB, cells were collected in 0.5 mL of PEB and then pelleted at 300 x g for 10 minutes at 4°C. Cells were then resuspended in 1x PBS at a concentration of 50,000 cells per 50 μL, with a viability ≥ 90% as determined by trypan blue staining and then used for downstream applications (see below).

### Single Cell RNA-Sequencing of Murine Brain Cells

scRNA-Seq libraries were generated using the 10x Chromium Single Cell 3′ v2 reagent kit, according to the manufacturer’s instructions, and were sequenced on an Illumina Nextseq500. Briefly, raw sequencing data were handled using the 10x Genomics Cell Ranger software (www.10xgenomics.com). Fastq files were mapped to the mm10 genome, and gene counts were quantified using the Cellranger count function. Subsequently, expression matrices from each experiment were merged and then imported into Seurat (version 4.0.4, https://satijalab.org/seurat/) for log normalization. Cells with a percentage of mitochondrial reads above 10%, and with less than 250 features, were filtered out. Batch effects were corrected by regressing out the number of mitochondrial read percentage using the vars.to.regress function. Doublet contamination was removed using DoubletFinder (version 2.0.3) (McGinnis et al., 2019). Sample integration was achieved using SCTransform (version 0.3.2) (Hafemeister and Satija, 2019) before running principal component analysis (PCA) was performed and significant principal components were used for graph-based clustering. UMAP was used for 2-dimensional visualization (https://github.com/lmcinnes/umap). Differential expression of genes per cluster was performed using the Wilcoxon rank sum test (FindMarkers function default). For pseudotemporal analysis, normalized data from endothelial cells were passed from Seurat to Monocle2 (Qiu et al., 2017a; Qiu et al., 2017b; Trapnell et al., 2014) (version 2.20.2). The Monocle2 BEAM statistical test was utilized to determine genes changing in a pseudo temporal manner. To identify transcription factors regulating the changes in gene expression across endothelial development, we use SCENIC (version 1.2.4) (Aibar et al., 2017). Both +/- 500 bp and +/- 10 kb around the murine TSS motif ranking databases were used for the analysis with default parameters. Genes that were co-regulated by two or more regulons were visualized using Cytoscape (version 3.8.2) (Shannon et al., 2003).

To identify receptor-ligand interactions, we subset the endothelial, pericyte, and microglia clusters from E9.5 and adult mice. The Wilcoxon signed ranked test was used to identify differentially expressed genes between timepoints in each cluster. Only genes present in at least 10% of cells, and with a log fold change above 0.25, were considered. We then use CCInx (version 0.5.1, (Ximerakis et al., 2019) to identify interaction between cell populations across time. Results can be accessed at the interactive shinyapp (https://mcantug.shinyapps.io/Endo_CCInxE9Ad/). Upstream regulation of differentially expressed genes in E9.5 and adult samples was analyzed and visualized by circus plot using NicheNET (version 1.0.0) (Browaeys et al., 2020) with default parameter settings. Only active ligands at the 95^th^ quantile was shown.

### OMNI-ATAC and RNA-Seq of Blood-Brain Barrier hCMEC/D3 Cells

Immortalized hCMEC/D3 (Millipore, SCC066) cells were grown to confluence using endothelial cell medium (ScienCell, #1001) on plates coated with Collagen Type I Rat Tail (Sigma-Aldrich, #C7661). Passages 4-6 were used for experiments. ATAC libraries were processed as previously described (Corces et al., 2017). The quality of purified DNA libraries was assessed using the Agilent High Sensitivity DNA kit (Agilent Technologies). Paired end, 2 x 75 bp sequencing was performed on an Illumina Nextseq 500 instrument. Reads were mapped to the GRCh38 version of the human genome using Bowtie2 with default paired-end settings (Langmead and Salzberg, 2012). Mitochondrial reads, reads with a MAPQ < 10, and reads which did not align to the reference genome were removed using Samtools (version 1.13) (Danecek et al., 2021). Duplicated reads were then removed with Picard MarkDuplicates (Institute, 2019). Peak calling was carried out with MACS2 (callpeak --nomodel –broad) (v2.2.7.1)(Zhang et al., 2008). Diffbind (version 3.2) (Ross-Innes et al., 2012; Stark R, 2011) was used to import peaksets (min.overlap= 0.66) into RStudio Server (version 1.4.1717, https://www.rstudio.com) using R (version 4.1 (Team, 4.1). dba.blacklist function was used to filter out peaks that overlap with the ENCODE blacklist. Consensus peaks were converted to mm10 using the LiftOver tool available from the UCSC Genome Browser (https://genome.ucsc.edu/cgi-bin/hgLiftOver). A region was considered conserved if a minimum 0.95 ratio of bases remapped to the murine genome. Selected regions were also examined using the ECR Browser (Ovcharenko et al., 2004) where the regions were analyzed using rVista 2.0 (Loots and Ovcharenko, 2004) to identify conserved transcription factor motifs. The TRANSFAC professional V10.2 vertebrate library was used with default parameters.

RNA was isolated using Trizol. Upon processing, RNA from all samples was thawed and following confirmation of integrity and concentration using a Bioanalyzer, 100 ng was used for low-input library preparation using the NEBNext Ultra II RNA Library Prep kit for Illumina. The libraries were then quantified and sequenced using an Illumina NovaSeq 6000 at a depth of 20 million reads per sample. Reads were first mapped to the human genome (GRCh38) using Salmon (version 1.5.1) (Patro et al., 2017). Transcript level quantification was then imported using txtimport (version 1.20.0) (Soneson et al., 2015) and analyzed using DESeq2 (Love et al., 2014). Genes were kept and considered actively expressed if they had more than 10 raw counts and >2 log2 fold change normalized expression.

### Statistics

Unless otherwise indicated, experiments were performed using a minimum of 2 independent biological replicates.

## Data availability

Datasets generated within this manuscript were deposited to the Gene Expression Omnibus, (GEO: GSE185345. Human dataset GEO: GSE187565).

## Acknowledgments

We thank Dr. Jason Fish for critical comments on the manuscript, and Ms. Karen Berman de Ruiz and Dr. Alexander Herman for assistance with mouse husbandry and organ isolation. Diagrams in some figures were created using Biorender.com.

## Author Contributions

M.C.G., M.C.H. and J.D.W. were responsible for the conception, design, execution, and interpretation of experiments. M.C.G. and J.D.W. wrote the original draft. G.L. was involved in the design, execution, and analysis of experiments. J.F.M. contributed reagents and resources, supervised M.C.H., interpreted experiments, and edited the manuscript. All authors revised the manuscript and consented to its contents.

---

**Supplemental Figure 1.**
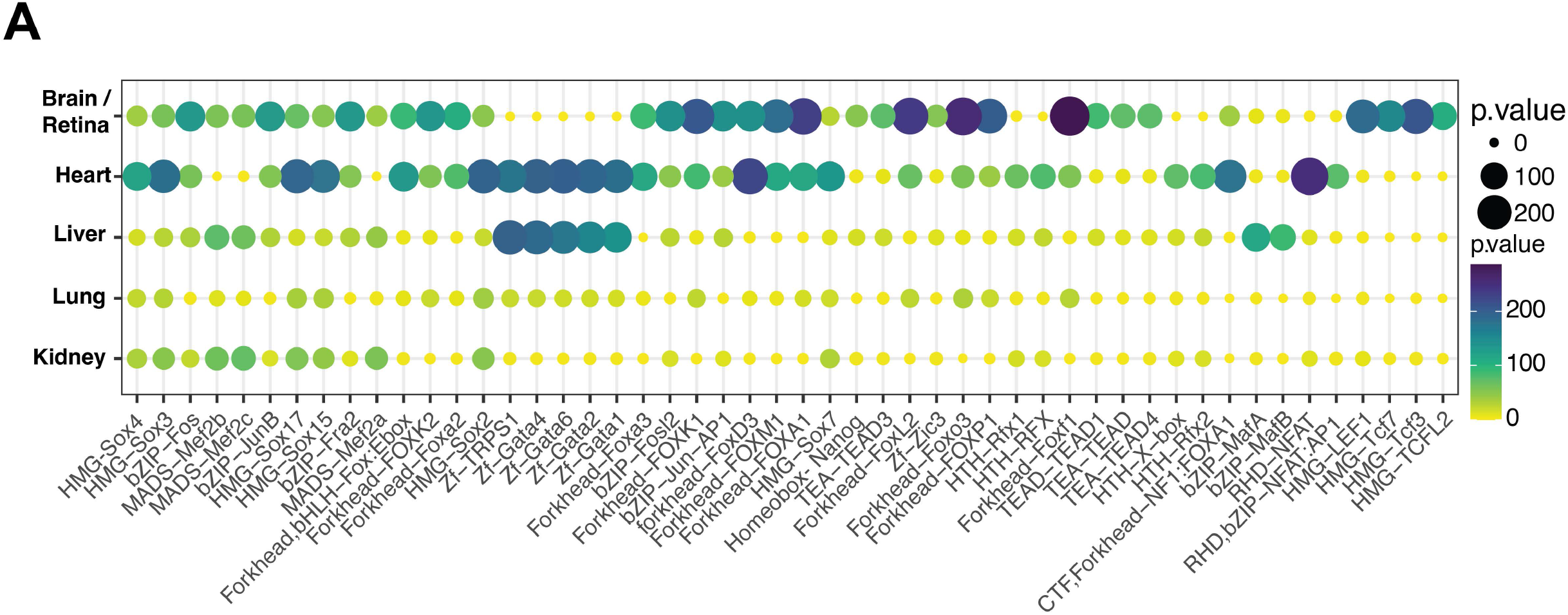
Top 50 Motifs Across all Organs. A) Enriched motifs identified by HOMER from all organs, with all timepoints condensed into one sample per organ. Size of the bubble and the color represent the p-value. The top 50 motifs are shown.

**Supplemental Figure 2.**
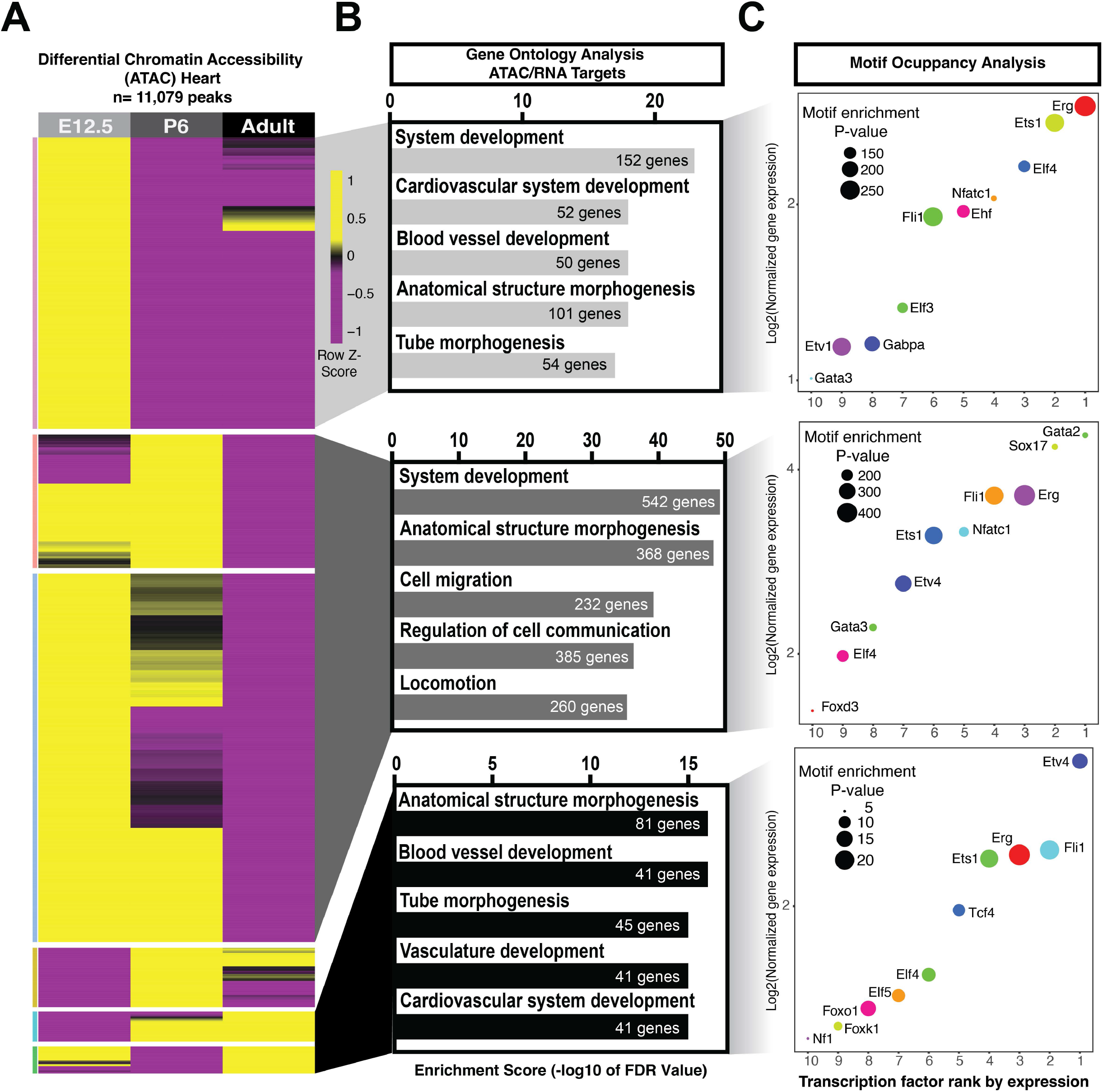
Chromatin Accessibility Changes Across Time in the Heart Endothelium. A) Differential chromatin accessibility determined by ATAC-Seq peaks in the heart endothelium (11,079 peaks) at E12.5, postnatal day 6 (P6) and adult (2-month-old) mice. B) Biological processes from expressed genes and with accessible chromatin in each timepoint. C) Top 10 transcription factor motifs ranked by gene expression for each age. Log2 expression over input indicated in the y-axis. Motif enrichment p-value is shown according to the dot size.

**Supplemental Figure 3.**
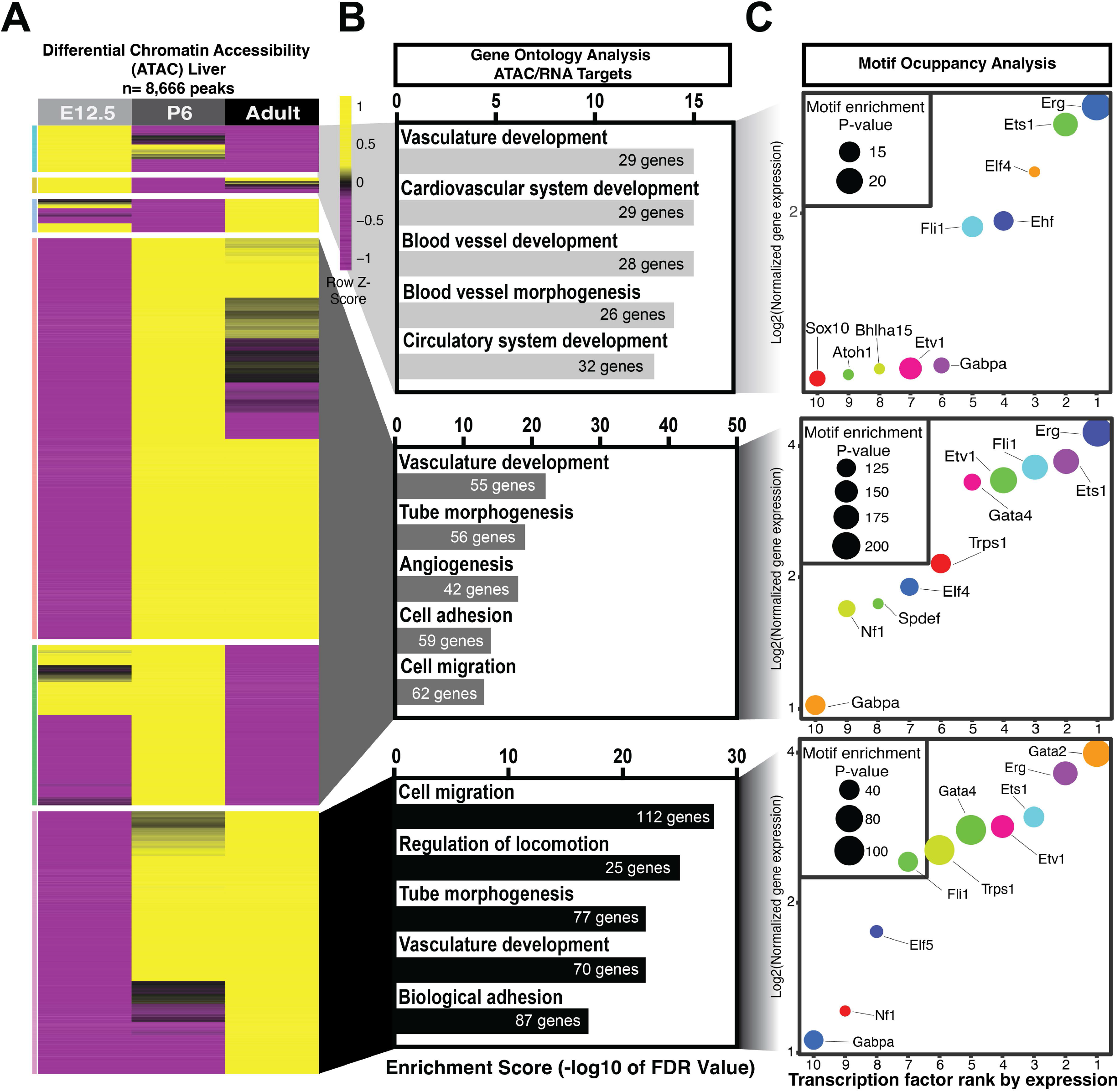
Chromatin Accessibility Changes Across Time in the Liver Endothelium. A) Differential chromatin accessibility determined by ATAC-Seq peaks in the liver endothelium (8,666 peaks) at E12.5, postnatal day 6 (P6) and adult (2-month-old) mice. B) Biological processes from expressed genes and with accessible chromatin in each timepoint. C) Top 10 transcription factor motifs ranked by gene expression for each age. Log2 expression over input indicated in the y-axis. Motif enrichment p-value is shown according to the dot size.

**Supplemental Figure 4.**
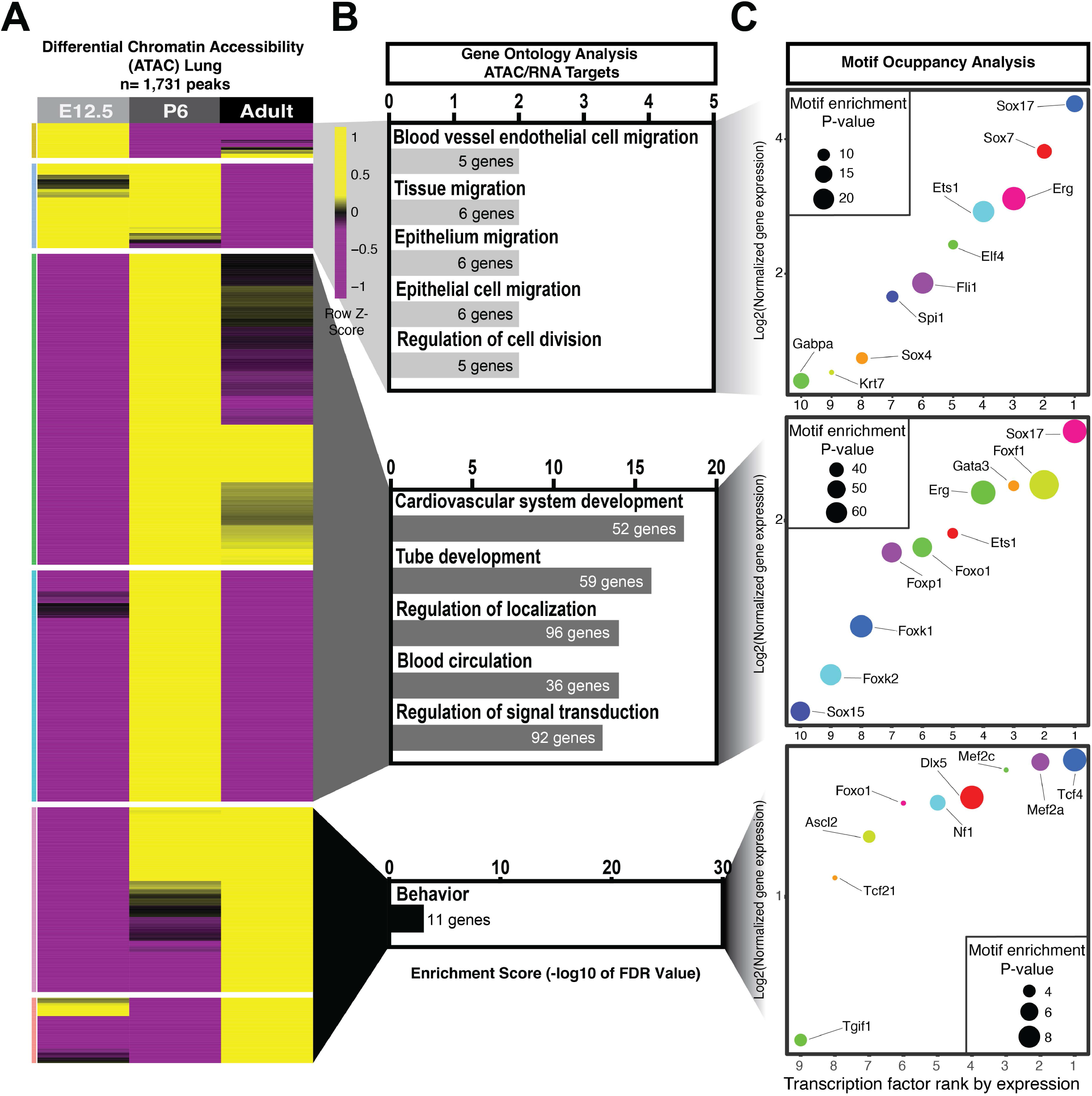
Chromatin Accessibility Changes Across Time in the Lung Endothelium. A) Differential chromatin accessibility determined by ATAC-Seq peaks in the lung endothelium (1,731 peaks) at E12.5, postnatal day 6 (P6) and adult (2-month-old) mice. B) Biological processes from expressed genes and with accessible chromatin in each timepoint. C) Top 10 transcription factor motifs ranked by gene expression for each age. Log2 expression over input indicated in the y-axis. Motif enrichment p-value is shown according to the dot size.

**Supplemental Figure 5.**
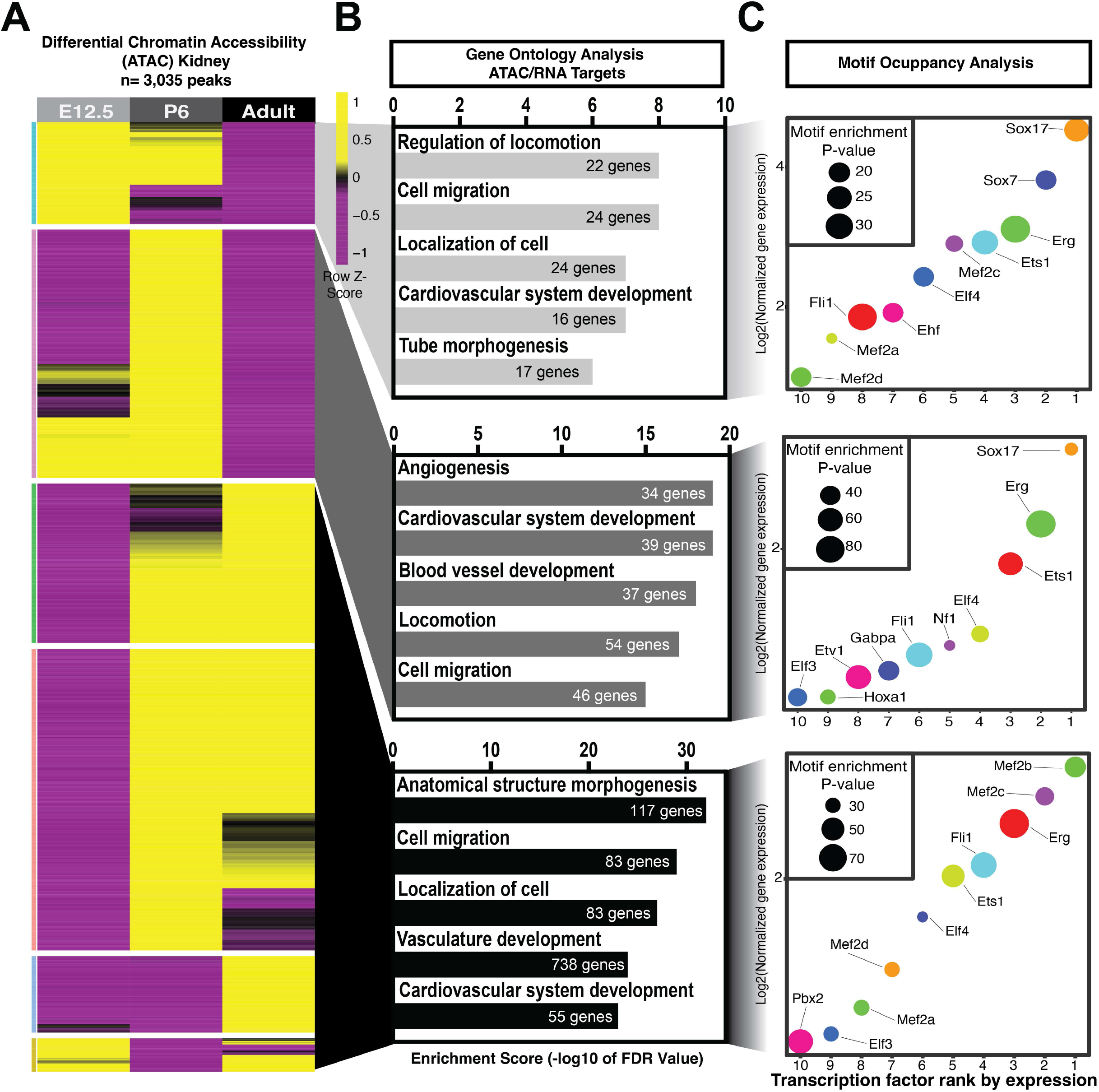
Chromatin Accessibility Changes Across Time in the Kidney Endothelium. A) Differential chromatin accessibility determined by ATAC-Seq peaks in the kidney endothelium (3,035 peaks) at E12.5, postnatal day 6 (P6) and adult (2-month-old) mice. B) Biological processes from expressed genes and with accessible chromatin in each timepoint. C) Top 10 transcription factor motifs ranked by gene expression for each age. Log2 expression over input indicated in the y-axis. Motif enrichment p-value is shown according to the dot size.

**Supplemental Figure 6.**
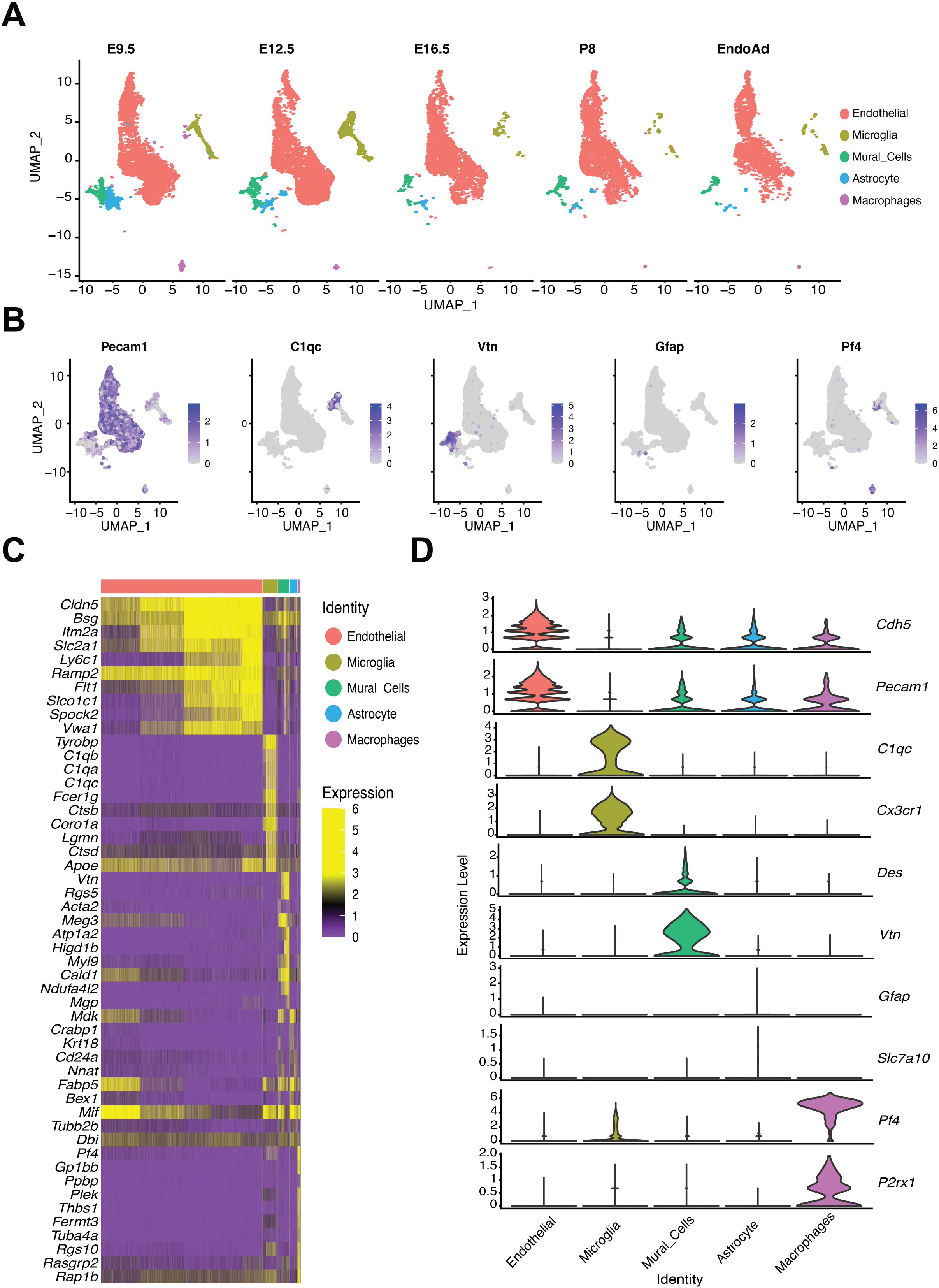
Classification of Major Cell Types Using Single Cell Sequencing. A) UMAP representation of different cell type clusters across timepoints. B) UMAP visualization of marker genes in selected clusters. C) Heatmap of the top 10 differentially expressed genes across cell types. D) Violin plots showing gene expression distribution of two canonical gene markers for each cell type.

**Supplemental Figure 7.**
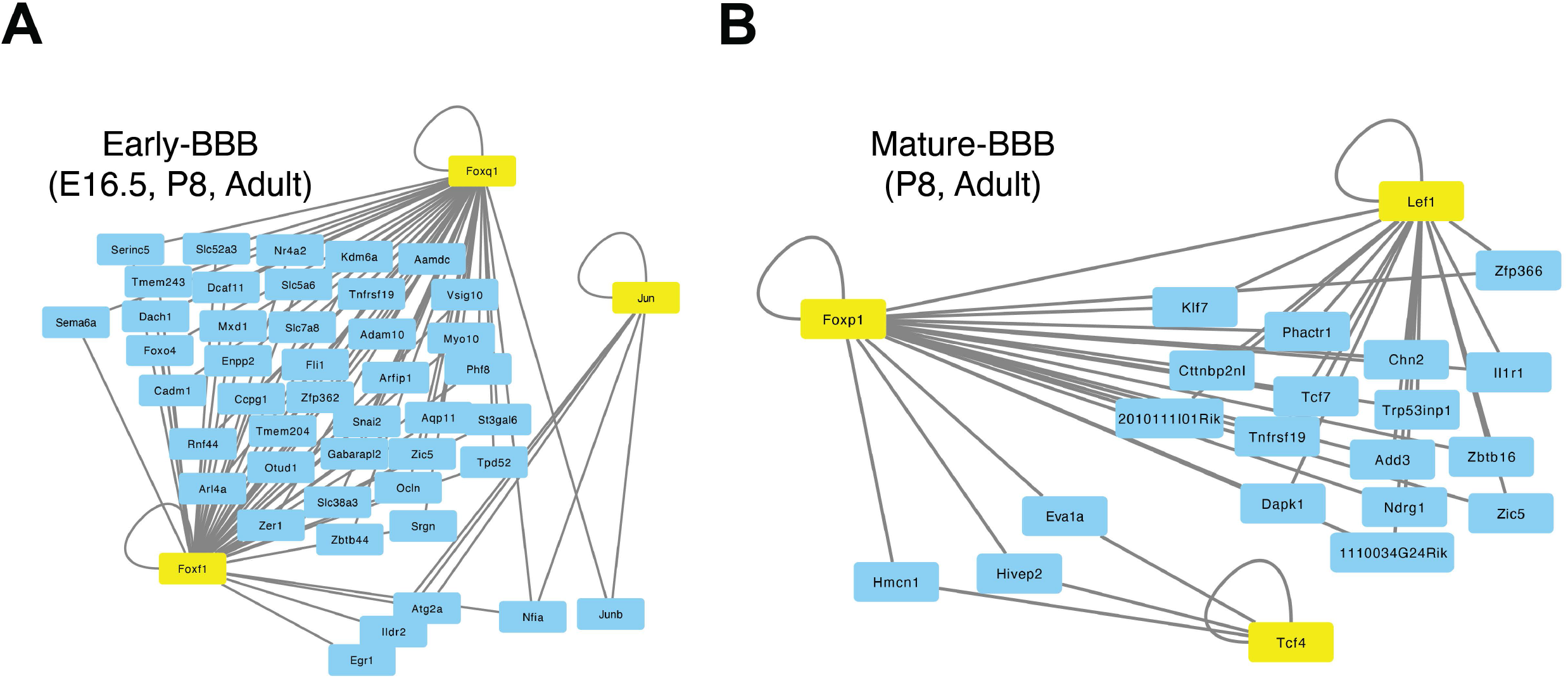
Common Target Genes in Active Regulons within the Developing and Mature Brain Endothelium. Interaction network constructed from the top 3 regulons, as determined by SCENIC, of the E16.5, P8 and adult (A) or P8 and adult only (B) brain endothelium. Genes regulated by 2 or more transcription factors are shown.

**Supplemental Figure 8.**
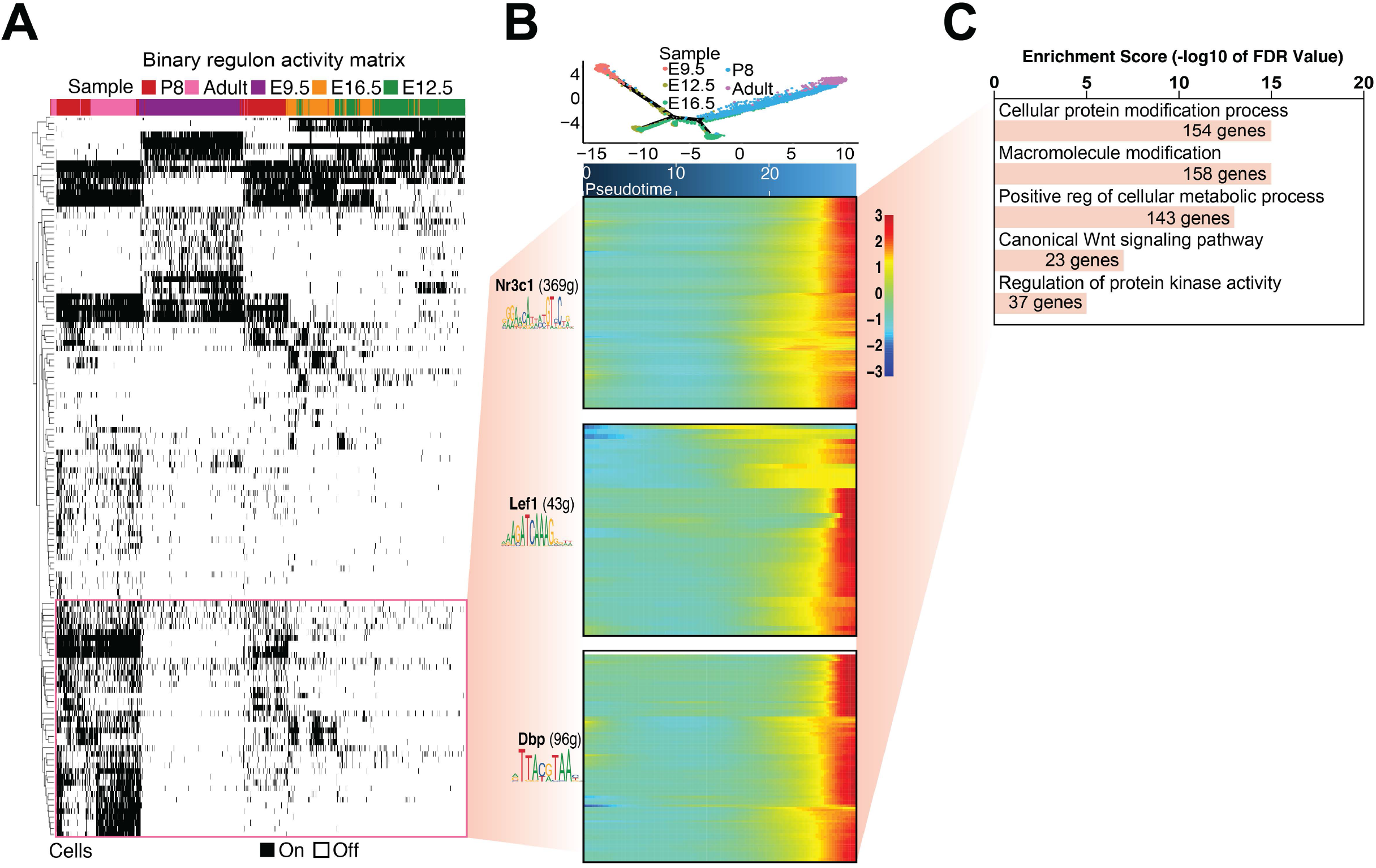
Mature BBB Regulon Activity Across Time and Gene Ontology Analysis. A) SCENIC binary activity heatmap representing active regulons in brain endothelial cells across all timepoints. Highlighted regulons are shown in panel B. B) Heatmaps show differentially active regulon target gene expression in the cerebral endothelium in P8 and adult. C) Selected GO biological processes derived from the target genes expressed by the three regulon clusters shown in panel B.

**Supplemental Figure 9.**
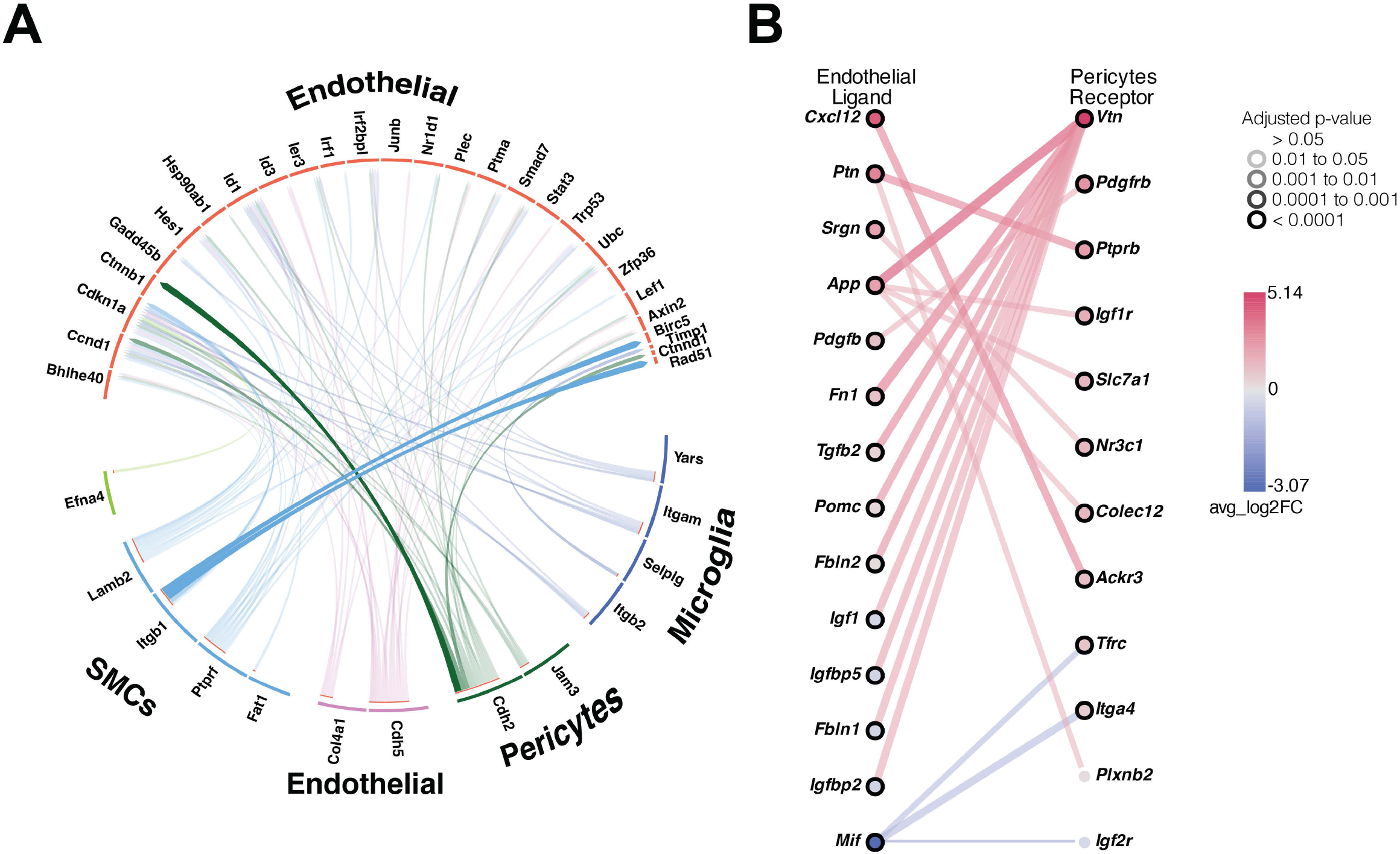
Cell to Cell Communication Changes in the Neurovascular Unit Over Time. A) Circos plot of differentially expressed ligands in non-EC cells within our dataset, as well as their target genes expressed in the CNS endothelium between E9.5 and Adult. F) Unbiased analysis of top predicted interactions of differentially expressed ligands and receptors between ECs and pericytes in E9.5 and adult using the Cell-Cell Interactions (CCInx).

**Supplemental Table 1. List of samples sequenced. Shared endothelial peaks across organs and timepoints. Gene Ontology (GO) terms and HOMER Motifs associated with shared peaks. Erg and Fli1 motif annotated peaks (associated with** Figure 2).

**Supplemental Table 2. Organ specific peaks and associated genes. Gene Ontology (GO) terms for each organ (associated with** Figure 3).

**Supplemental Table 3. Lef1, Nfat, Gata4, Foxo3 and Hoxc9 annotated target peaks (associated with** Figure 3).

**Supplemental Table 4. E12.5, P6 and adult brain peaks, annotated target genes and Gene Ontology (GO) terms associated with it (associated with** Figure 4).

**Supplemental Table 5. Differentially expressed genes in annotated single cell clusters. Differentially expressed genes in endothelial cells across timepoints (associated with** Figure 5).

**Supplemental Table 6. Target genes of selected regulons and Gene Ontology (GO) terms divided by developmental stage (associated with** Figure 6).

**Supplemental Table 7. Conserved ATAC regions between hCMEC/D3 cells and adult mouse brain endothelium with HOMER motif analysis.**

## SUPPLEMENTAL MATERIALS AND METHODS

**Table S1:**
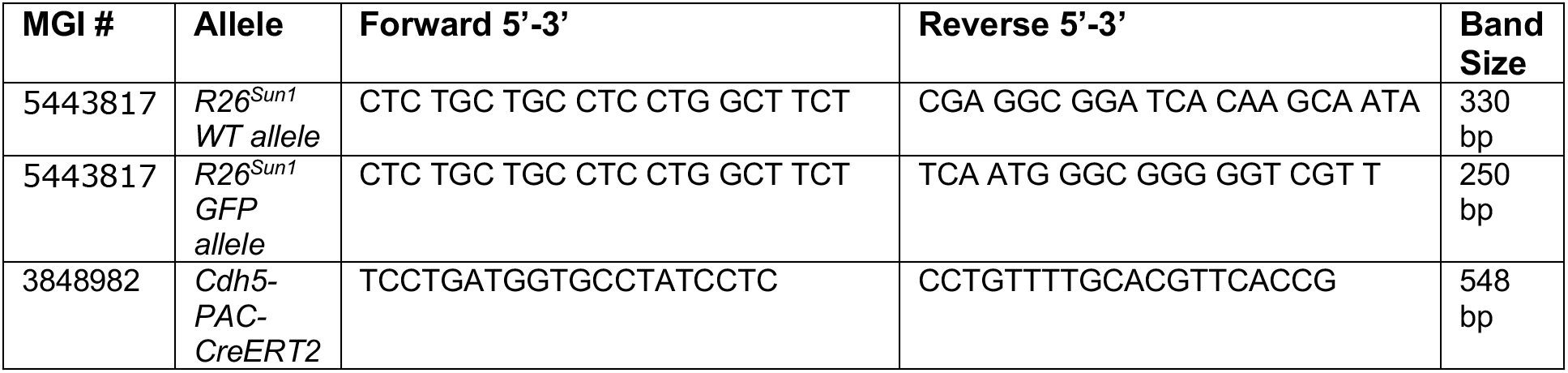
Primers used for murine genotyping.

